# Mitochondrial bioenergetics stimulates autophagy for pathological tau clearance in tauopathy neurons

**DOI:** 10.1101/2024.02.12.579959

**Authors:** Nuo Jia, Dhasarathan Ganesan, Hongyuan Guan, Yu Young Jeong, Sinsuk Han, Marialaina Nissenbaum, Alexander W. Kusnecov, Qian Cai

**Affiliations:** Division of Life Sciences, Department of Cell Biology and Neuroscience and; Department of Psychology, School of Arts and Sciences, Rutgers, The State University of New Jersey, Piscataway, NJ 08854, USA

## Abstract

Hyperphosphorylation and aggregation of microtubule-associated tau is a pathogenic hallmark of tauopathies and a defining feature of Alzheimer’s disease (AD). Pathological tau is targeted by autophagy for clearance, but autophagy dysfunction is indicated in tauopathy. While mitochondrial bioenergetic failure has been shown to precede the development of tau pathology, it is unclear whether energy metabolism deficiency is involved in tauopathy-related autophagy defects. Here, we reveal that stimulation of anaplerotic metabolism restores defective oxidative phosphorylation (OXPHOS) in tauopathy which, strikingly, leads to enhanced autophagy and pronounced tau clearance. OXPHOS-induced autophagy is attributed to increased ATP-dependent phosphatidylethanolamine biosynthesis in mitochondria. Excitingly, early bioenergetic stimulation boosts autophagy activity and reduces tau pathology, thereby counteracting memory impairment in tauopathy mice. Taken together, our study sheds light on a pivotal role of bioenergetic dysfunction in tauopathy-linked autophagy defects and suggests a new therapeutic strategy to prevent toxic tau buildup in AD and other tauopathies.

## Introduction

Alzheimer’s disease (AD) is pathologically characterized by the presence of extracellular amyloid plaques consisting of amyloid β (Aβ) peptides and intracellular neurofibrillary tangles (NFTs) composed of hyperphosphorylated tau (phospho-tau) in diseased brains [1,2]. In AD, tau pathology correlates with dementia better than amyloid plaques [3]. Tau is encoded by the *MAPT* (microtubule-associated protein tau) gene. Mutations in the *MAPT* gene cause a subtype of frontotemporal dementia (FTD) and overexpression of FTD-associated mutant *MAPT* genes in mice results in NFT development and neurodegeneration [4-6], establishing a central role of pathological tau in tauopathy. Thus, a deeper understanding of the pathogenic mechanisms of tau is of great importance in developing tau-targeted therapies to treat AD and other tauopathies. Phospho-tau is degraded within lysosomes after being targeted by autophagy, a key cellular quality-control mechanism for the maintenance of neuronal homeostasis and brain health [7-14]. Studies by us and others have uncovered that autophagosomes/autophagic vacuoles (AVs) are primarily formed in axons and synapses, and that nascent AVs loaded with engulfed cargos move in a predominantly retrograde direction toward the soma of neurons for lysosomal degradation [15-21]. Previous work indicated the perturbation of autophagy function in both tauopathy patient brains and cellular models [22-25]. However, the underlying mechanism remains largely unknown.

The formation of an AV is a hallmark of autophagy, which is initiated with the generation and elongation of a pre-autophagosomal membrane (i.e., phagophore or isolation membrane) that is finally enclosed to form an AV [11,12,26,27]. The life cycle of AVs requires the organized accretion of phospholipids at the nucleation site and during AV membrane elongation, cargo engulfment, and resolution through fusion with lysosomes, making these phospholipids a limiting factor for autophagy activity [26,28,29]. The origin of AVs and the lipid sources for AV biogenesis have been under extensive scrutiny only in recent years. Mounting evidence indicates that phosphatidylethanolamine (PE), a major component of the AV membrane, is critically involved in autophagy initiation and AV membrane elongation [30]. PE functions as an anchor to enable the association of autophagy-related protein LC3 (Atg8) with the AV membrane, which is crucial for AV biogenesis *de novo* [31-33]. In mammalian cells, PE is mainly synthesized via the CDP-ethanolamine (ETN) (Kennedy) pathway in the endoplasmic reticulum (ER), and via phosphatidylserine (PS) decarboxylase (PSD) in mitochondria [34-36]. In the ER CDP-ETN pathway, ETN is converted to PE by the sequential actions of ETN kinase, CTP: phosphoethanolamine cytidylyltransferase and finally choline/ethanolaminephosphotransferase. In the PSD pathway, PS is decarboxylated by the enzyme PSD to generate PE in mitochondria. It is also important to note that PE metabolism in both the ER and mitochondria is an energy-requiring process that is fueled by adenosine triphosphate (ATP) [37,38]. This makes PE biosynthesis a potentially critical node connecting cellular bioenergetic status and autophagy. However, it is still unclear which energy source powers PE metabolism within cells, and whether ATP-dependent PE biosynthesis in the ER, mitochondria, or both, is critical in autophagy functionality for the prevention of toxic tau accumulation in disease neurons.

Mitochondria are cellular energy powerhouses that supply the majority of the cell’s energy in the form of ATP, which is produced through oxidative phosphorylation (OXPHOS) and mediated by the electron transport chain (ETC) system embedded in the inner mitochondrial membrane (IMM). ATP supplied by mitochondria powers a variety of activities essential for neuronal function and survival. Mitochondrial dysfunction in the nervous system is a central concern during aging and has been associated with major incapacitating neurodegenerative disorders, including the highly prevalent AD [39-42]. Volumes of evidence suggest that mitochondrial bioenergetic disruption constitutes one of the earliest and most notable features of AD before any histopathological or clinical abnormalities [43-45]. We and others have revealed that mitochondrial bioenergetic failure and energy crisis are early events preceding the development of tau pathology [46-49]. However, it is yet to be determined whether ATP supply through OXPHOS is crucial for PE metabolism, whether mitochondrial energy metabolism deficiency disrupts the ATP-requiring biosynthesis of PE for autophagy, and whether early tauopathy-associated OXPHOS deficits play a role in the pathogenesis of autophagy defects that result in pathological tau buildup.

The present study has revealed that, in tauopathy neurons, stimulation of anaplerotic metabolism to boost OXPHOS activity enhances PE biosynthesis in mitochondria, but not in the ER. This induces a robust increase in autophagy and leads to drastic pathological tau clearance. Excitingly, early enhancement of mitochondrial bioenergetics rescues autophagy deficits, attenuates tau pathology, and protects against neurodegeneration and cognitive impairment in tauopathy mice. Therefore, our findings highlight a novel role of mitochondrial bioenergetics in the regulation of neuronal autophagy and provide new insights into the potential involvement of mitochondrial energy metabolism dysfunction in autophagy failure and pathological tau accumulation in the context of tauopathy. In addition, the current work strengthens our understanding of tauopathy and other neurodegenerative diseases as well as aging associated with bioenergetic deficiency and autophagy defects. Our work suggests the stimulation of anaplerotic metabolism as a new, potentially therapeutic strategy aiming to enhance autophagy functionality and thus promote toxic tau clearance in tauopathies, including AD.

## Results

### Stimulation of anaplerotic metabolism corrects mitochondrial bioenergetic deficits in tauopathy neurons

Bioenergetic deficits are an early feature of tauopathy [46-49]. Therefore, we sought to address whether such deficits could be reversed by stimulation of OXPHOS activity. We induced anaplerotic metabolism in primary cortical neuron cultures derived from non-Tg or tauP301S Tg (PS19) mouse brains by switching from the media containing glucose but no glutamine to glucose-free media supplemented with glutamine (Gln). Glutamine is an anaplerotic amino acid that stimulates OXPHOS by being converted into α-ketoglutarate, an intermediate of the tricarboxylic acid (TCA) cycle [50-53]. We and others have demonstrated that cells under glutamine-supplemented conditions exhibited a significant increase in the basal oxygen consumption rate (OCR) compared to the cells grown in the presence of glucose only [54-58]. The basal OCR of the cells reached a saturated level with the concentration of supplemented glutamine between 8 mM and 12 mM [56]. Thus, we applied 10 mM glutamine to PS19 neuron cultures for 24 hours. We first assessed the effects of glutamine supplementation on OXPHOS activity by examining ATP levels in mitochondria in the soma and the axons of neurons. We utilized mitAT1.03, a genetically encoded fluorescence resonance energy transfer (FRET)-based ATP indicator, which is localized to the mitochondrial matrix through the N terminus of AT1.03 fusion to a duplex of the mitochondrial targeting signal of Cytochrome c oxidase subunit VIII [59]. For the AT1.03 probe, the ε subunit of *Bacillus subtilis* FoF1-ATP synthase is connected to YFP and CFP. The ATP-bound form of the ε subunit of *Bacillus subtilis* FoF1-ATP synthase increases FRET efficiency, as reflected by an enhanced FRET signal (YFP/CFP emission ratio) which indicates an increase in ATP levels. Thus, mitAT1.03 and AT1.03 allow for monitoring of ATP levels in the mitochondria and the cytoplasm, respectively. Relative to those in non-Tg neurons, we observed a significant reduction in YFP/CFP ratios of these mitochondria in the soma and the axons of PS19 neurons (p < 0.0001) (Figure 1A and B), suggesting defective mitochondrial bioenergetics in these neurons. We further determined whether elevated OXPHOS activity increases ATP levels in the cytoplasm by using AT1.03. In glutamine-free media, PS19 neurons exhibited markedly reduced cellular ATP levels, as evidenced by decreased YFP/CFP ratios compared to those in non-Tg neurons (Figure S1A and B). Such a defect was corrected in PS19 neurons under glutamine oxidation, which showed higher YFP/CFP ratios in both the soma and the axons than those in control PS19 neurons in the absence of glutamine (p < 0.0001) (Figure S1A and B). These observations indicate that the anaplerotic stimulation of OXPHOS rescues energy deficits in tauopathy neurons. Notably, we also found a significant increase in the density of synaptophysin (SYP)-indicated presynaptic terminals accompanied by elevated presynaptic mitochondrial contents in the axons of glutamine-treated PS19 neurons (p < 0.0001) (Figure 1C and D). In OXPHOS-stimulated PS19 neurons, we did not detect significant changes in Syntaxin 1 (STX1), a neuronal marker, as well as mitochondrial proteins. Of note, both biochemical and imaging data showed an increase in SYP (Figure 1C-F), suggesting that anaplerotic stimulation results in beneficial effects at tauopathy synapses. Furthermore, these glutamine-treated PS19 neurons displayed a robust reduction in the levels of tau, visualized by Tau5 and PHF1 antibodies that recognize total tau and phospho-tau, respectively (Figure 1E and F). This interesting finding supports us in further defining the mechanism underlying high OXPHOS activity-induced tau reduction in tauopathy neurons.

**Figure 1.**
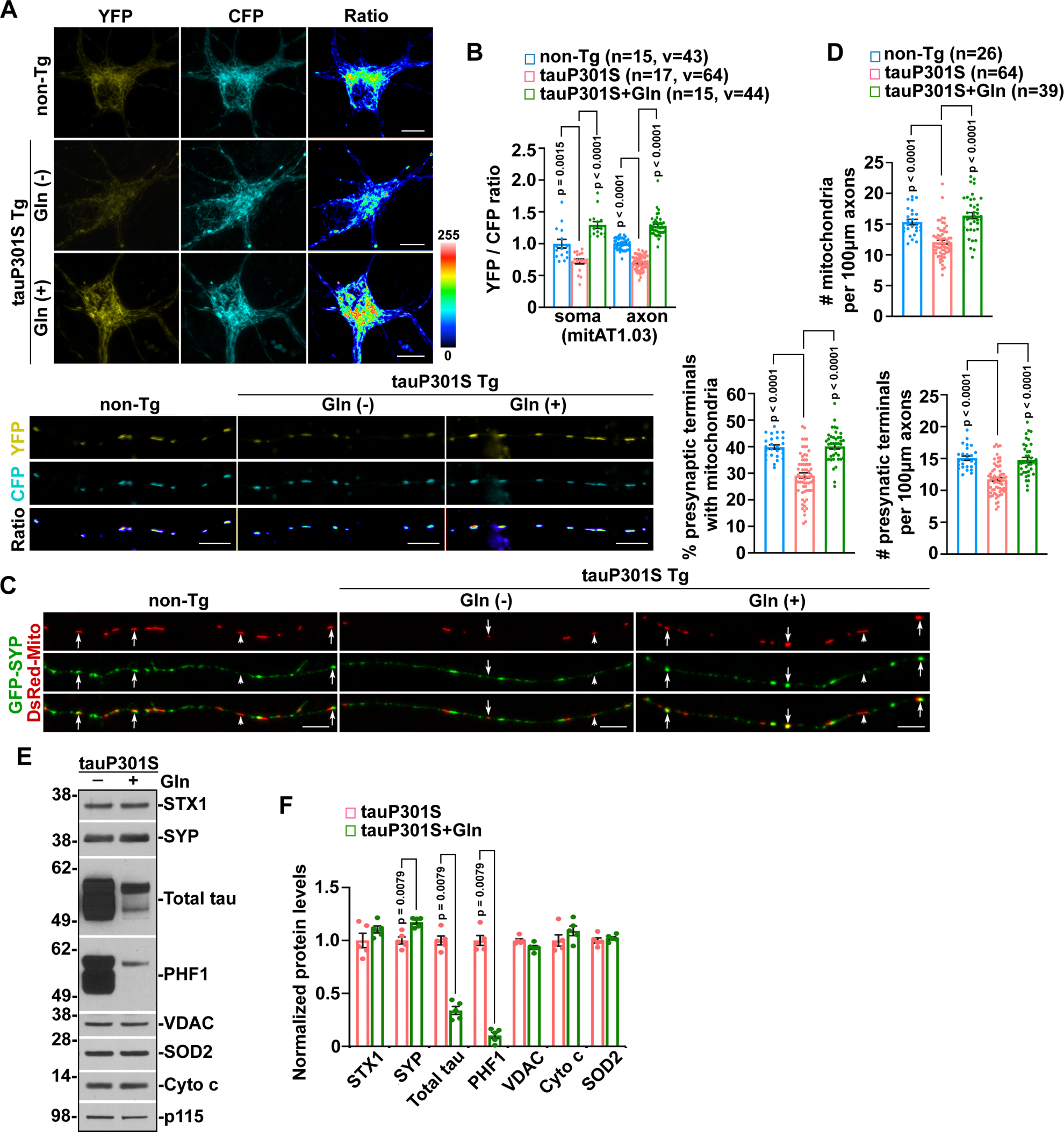
Stimulation of anaplerotic metabolism elevates oxidative phosphorylation (OXPHOS) activity in tauopathy neurons. (A-B) Representative images (A) and quantitative analysis (B) of mitochondrial ATP levels indicated by mitAT1.03 in DIV10-12 primary cortical neurons cultured from non-transgenic (Tg) and tauP301S Tg (PS19) mouse brains in the presence and absence of 24-hour glutamine (10 mM) treatment. The YFP/CFP ratios of the somatic and axonal mitochondria in PS19 neurons with and without glutamine supplementation were normalized to those in control non-Tg neurons, respectively. Gln: glutamine. (C-D) Representative images (C) and quantitative analysis (D) of the densities of mitochondria, presynaptic terminals with mitochondria, and presynaptic terminals in the axons of non-Tg neurons or PS19 neurons at DIV12-14 with and without 24-hour 10 mM glutamine supplementation. Arrows: synaptic mitochondria; arrowheads: non-synaptic mitochondria SYP: synaptophysin. (E-F) Representative blots (E) and quantitative analysis (F) of protein levels in DIV12-14 PS19 neurons in the presence and absence of 24-hour 10 mM glutamine treatment. Total tau and phospho-tau were visualized by Tau5 and PHF1 antibodies, respectively. The protein levels were normalized to the loading control p115 and to those of control PS19 neurons with no glutamine supplementation, respectively. STX1: Syntaxin 1; Cyto c: Cytochrome c. p115, a Golgi protein. Data were collected from a total number of neuronal soma (n) or axons (v) indicated in parentheses (B, D) from five independent experimental repeats or were quantified from five independent repeats (F). Data were expressed as the mean ± SEM with dots as individual values and analyzed by one-way ANOVA with Bonferroni’s correction (B and D) or Mann-Whitney U test (F). Scale bars: 10 μm.

### OXPHOS enhancement leads to pronounced pathological tau clearance by increasing AV biogenesis in tauopathy neurons

We first determined whether OXPHOS-stimulated tau decrease is attributed to elevated tau turnover in PS19 neurons. Indeed, in the presence of cycloheximide (CHX) which blocks protein synthesis, the levels of total tau and phospho-tau species visualized by Tau5, PHF1, and AT8 antibodies showed marked reductions in PS19 neurons supplemented with glutamine (Figure 2A and B), suggesting increased tau turnover under glutamine metabolism. Given that pathological tau is targeted by autophagy for clearance [7-10], we also assessed the levels of LC3-II, a marker of autophagy [60]; sequestosome 1 (p62/SQSTM1), an adaptor protein that selects ubiquitinated proteins for autophagy [61]; and LAMP1, a lysosomal protein. Strikingly, in OXPHOS-enhanced PS19 neurons, we observed drastically increased levels of LC3-II and LAMP1 along with decreased p62 (Figure 2A and B). Moreover, OXPHOS stimulation-induced reduction of tau, particularly oligomeric tau, was suppressed in PS19 neurons in the presence of lysosomal inhibitors (LIs) (monomeric tau: p < 0.0001; oligomeric tau: p < 0.0001) (Figure 2C and D), indicating that the tau clearance is lysosome-dependent. These findings support the possibility that glutamine-enhanced OXPHOS promotes tau clearance in PS19 neurons by elevating autophagy activity. To confirm the effects of anaplerotic stimulation on autophagy and tau clearance in PS19 neurons, we applied another anaplerotic amino acid aspartate. Similar to the effects of glutamine supplementation, we observed increased LC3-II and decreased p62 along with decreases in the levels of total tau and phospho-tau in PS19 neurons in the presence of aspartate (AT8: p = 0.0004; PHF1: p = 0.0002; Tau5: p < 0.0001; LC3-II: p = 0.0012; p62: p < 0.0001) (Figure S2A and B). Collectively, these results indicate that OXPHOS enhancement boosts autophagy functionality for the removal of pathological tau species.

**Figure 2.**
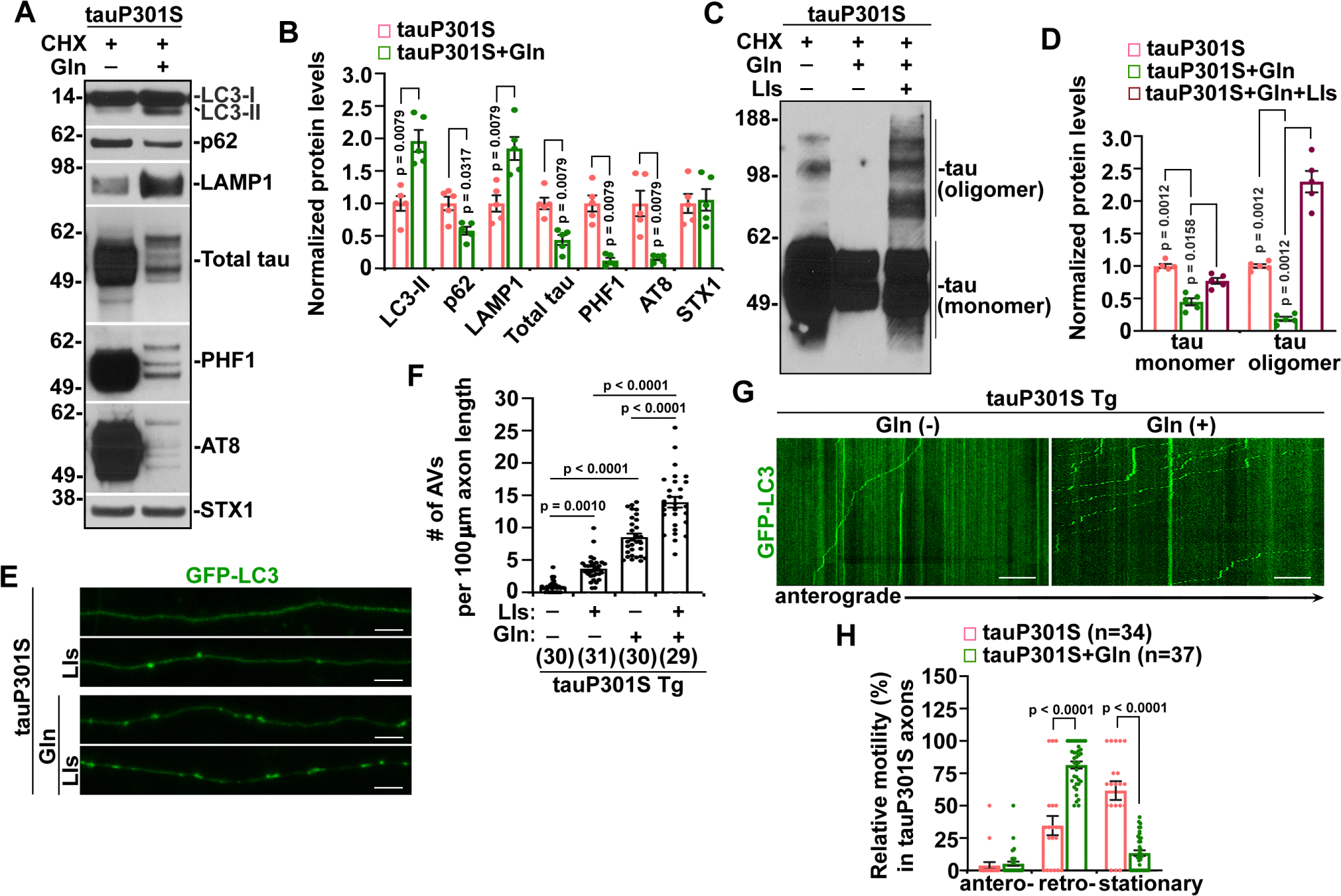
Anaplerotic stimulation of OXPHOS in tauopathy neurons enhances tau clearance by increasing autophagic flux. (A-B) Representative blots (A) and quantitative analysis (B) of autophagy and tau protein levels in PS19 neurons at DIV12-14 treated 24 hours with 10 μg/ml CHX or CHX and 10 mM glutamine. The protein levels were normalized to STX1 and to those of control PS19 neurons in the absence of glutamine, respectively. CHX: cycloheximide. (C-D) Representative blots (C) and quantitative analysis (D) of monomeric and oligomeric tau levels revealed by a native gel in DIV12-14 PS19 neurons incubated 24 hours with CHX, CHX and glutamine, or CHX, glutamine, and LIs (20 μM Pepstatin A and 20 μM E64D). The protein levels were normalized to those of PS19 neurons with CHX alone. LIs: lysosomal inhibitors. (E-F) Representative images (E) and quantitative analysis (F) of autophagic flux in the axons of DIV10-12 PS19 neurons in the presence and absence of LIs, glutamine, or glutamine and LIs for 24 hours. The data were quantified and expressed as the number of GFP-LC3-marked AVs per 100 μm axonal length in PS19 neurons. AV: autophagosome/autophagic vacuole. (G-H) Representative kymographs (G) and quantitative analysis (H) of AV motility in the axons of PS19 neurons with and without 24-hour glutamine treatment. Data were quantified from five independent repeats (B and D) or were collected from the total number of neurons (n) as indicated in parentheses (F and H) from more than three independent experiments. Data were expressed as the mean ± SEM and analyzed by Mann-Whitney U test (B), Kruskal-Wallis test with Dun’s multiple comparison post hoc test (D), one-way ANOVA with Bonferroni’s correction (F), or two-sided unpaired Student’s *t*-test (H). Scale bars: 10 μm.

Next, we sought to address whether autophagic flux is increased in bioenergetically enhanced PS19 neurons. We and others have established that AVs are primarily generated in the axons and synapses of neurons [15-21]. We thus assessed the density of autophagosomes/autophagic vacuoles (AVs) in the distal axons of PS19 neurons with and without glutamine treatment. Compared to PS19 neurons under basal conditions, we indeed found a drastic increase in the number of AVs in PS19 axons supplemented with glutamine, which became even greater when lysosomal inhibitors (LIs) were present (p < 0.0001) (Figure 2E and F). The soma of PS19 neurons showed similar increases in AV numbers under glutamine metabolism (p = 0.0002) (Figure S2C and D). These observations suggest that stimulation of anaplerotic metabolism elevates AV biogenesis in PS19 neurons. We further carried out time-lapse confocal imaging in live PS19 axons and demonstrated that OXPHOS stimulation triggered robust AV formation and that newly formed AVs moved in a predominantly retrograde direction toward the soma (Figure 2G and H). Moreover, similar results can be detected in the axons of non-Tg neurons supplemented with glutamine (AVs: p < 0.0001) (Figure S2E-H).

### OXPHOS-stimulated autophagy is mediated by mitochondrial PE biosynthesis in tauopathy neurons

We next determined the source of PE, a major component of the AV membrane [30], that supports OXPHOS-stimulated AV biogenesis in tauopathy neurons. Previous studies reported a key role of the ER in basal autophagy in dorsal root ganglion (DRG) neurons [16]. We thus applied meclizine, a specific inhibitor of the rate-limiting step in the ER CDP-ETN pathway [62]. Surprisingly, treatment of PS19 neurons with 5 μM meclizine for 24 hours showed no detectable effects on anaplerotic metabolism-stimulated autophagy (LC3-II: p = 0.0021; p62: p = 0.0029) and tau clearance (total tau: p = 0.0021; PHF1: p = 0.0024) (Figure S3A and B). Moreover, our imaging studies showed that ETN, which favors ER PE biosynthesis, increased the number of AVs in PS19 axons in the absence of glutamine, whereas ETN-elevated AV formation was blocked by meclizine, which demonstrated meclizine specificity (Figure S3C and D). In addition, like meclizine, ETN also had no impact on autophagy stimulated by OXPHOS in PS19 axons (Figure S3C and D). These results suggest that PE biosynthesis in the ER is not involved in anaplerosis-stimulated autophagy, which supports us in testing whether mitochondria-supplied PE plays a major role. We treated PS19 neurons with hydroxylamine, an irreversible inhibitor of PSD in mitochondria, at 0.1 mM dose for 6 hours which was previously shown to not result in signs of apoptosis [38]. Interestingly, compared to those in PS19 neurons supplemented with glutamine only, glutamine-enhanced autophagy activity was abrogated when hydroxylamine was present, as evidenced by halted LC3-II increase and p62 decrease (LC3-II: p = 0.0029; p62: p = 0.0037) accompanied by a blockage of tau reduction (total tau: p = 0.0024; PHF1: p = 0.0029) (Figure 3A and B). In line with these biochemical data, our imaging studies further demonstrated that glutamine-stimulated AV biogenesis was abolished in PS19 axons in the presence of hydroxylamine but not meclizine (p < 0.0001) (Figure 3C and D). PSD RNAi showed similar suppression of AV biogenesis in glutamine-supplemented PS19 axons, but not in control PS19 axons (p < 0.0001) (Figure S3E and F). Combined, these results support the view that OXPHOS-enhanced autophagy and tau clearance rely on PE synthesized in mitochondria, but not in the ER in tauopathy neurons.

**Figure 3.**
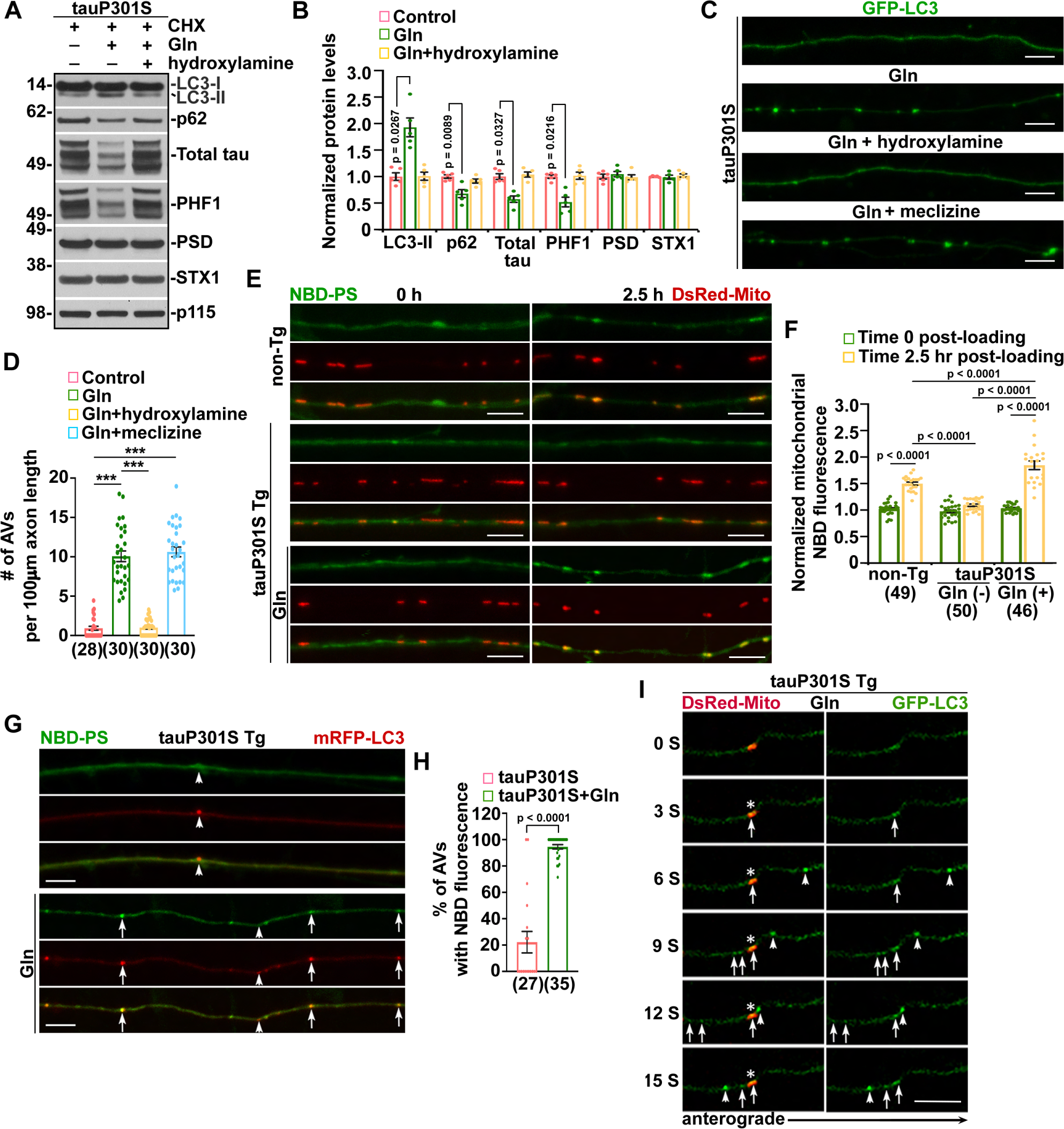
OXPHOS-enhanced autophagy is mediated by mitochondrial phosphatidylethanolamine (PE) biosynthesis in tauopathy neurons. (A-B) Representative blots (A) and quantitative analysis (B) of autophagy and tau protein levels in DIV12-14 PS19 neurons treated for 6 hours with 10 μg/ml CHX, CHX and 10 mM glutamine, or CHX, 10 mM glutamine, and 0.1 mM hydroxylamine. The protein levels were normalized to the loading control p115 and to those of control PS19 neurons with CHX alone, respectively. Data were quantified from five independent repeats. p115: a Golgi marker. (C-D) Representative images (C) and quantitative analysis (D) of autophagic flux in the axons of DIV10-12 PS19 neurons treated with 0.1 mM hydroxylamine for 6 hours or 5 μM meclizine for 24 hours in the presence or absence of 10 mM glutamine. The data were quantified and expressed as the number of GFP-LC3-labeled AVs per 100 μm axonal length in PS19 neurons. (E-F) Representative images (E) and quantitative analysis (F) of PE metabolism in mitochondria in the axons of DIV10-12 non-Tg neurons and PS19 neurons 2.5 hours following the loading of NBD-PS in the presence and absence of glutamine. The data were quantified and expressed as mitochondrial NBD-PE fluorescence that was normalized to non-mitochondrial NBD background signal from the same axons of non-Tg and PS19 neurons at time 0 and 2.5 hours, respectively. (G-H) NBD-labeling of AVs in the axons of DIV10-12 PS19 neurons 3 hours after NBD-PS loading with and without 3 hours glutamine preincubation. The data were quantified and expressed as the percentage of mRFP-LC3-indicated AVs positive for the NBD fluorescence in the axons of PS19 neurons with and without glutamine, respectively. Arrows: NBD-positive AVs; arrowheads: NBD-negative AVs. (I) Time-lapse imaging analysis of MDAV dynamics in PS19 axons administered with glutamine. Asterisks: DsRed-Mito-labeled mitochondria; arrows: MDAVs; arrowheads: a GFP-LC3-marked AV passing by the axon. MDAV: mitochondria-derived AV. Imaging data were quantified from the total number of neurons (n) as indicated in parentheses (D, F, and H) from more than three independent experiments. Data were expressed as the mean ± SEM and analyzed by Kruskal-Wallis test with Dun’s multiple comparison post hoc test (B), one-way ANOVA with Bonferroni’s correction (D and F), or two-sided unpaired Student’s *t*-test (H). Scale bars: 10 μm.

To confirm that OXPHOS-induced autophagy requires PE made in mitochondria, we directly determined the effects of anaplerotic stimulation of OXPHOS on mitochondrial PE metabolism in live tauopathy neurons by monitoring the conversion of the fluorescent analog of PS—18:1-06:0 N-[7-Nitrobenz-2-oxa-1,3-diazol-4-yl] phosphatidylserine (NBD-PS) to NBD-PE, which occurs only in mitochondria through the PSD pathway. Monitoring the accumulation and distribution of NBD-PE fluorescent signals in mitochondria allows for the assessment of mitochondrial PE metabolism and biosynthesis [63-66]. We found that the NBD-PE fluorescence signal in mitochondria reached its peak around 2.5 hours post-loading in control non-Tg axons. Importantly, the peak NBD-PE signal in the mitochondria of control PS19 axons was significantly decreased, which was reversed upon OXPHOS stimulation (p < 0.0001) (Figure 3E and F). We have shown bioenergetic deficits of the mitochondria in control PS19 axons in the absence of glutamine (Figures 1A and B, S1A and B). Thus, these results suggest that mitochondrial PE metabolism is impaired in PS19 neurons with bioenergetic deficits and that high OXPHOS activity stimulated by glutamine can rescue such defects. Furthermore, we assessed whether the mitochondria-synthesized PE is incorporated into nascent AVs in glutamine-treated PS19 axons. Indeed, 3 hours after the PS19 neuron culture was exposed to NBD-PS, most mRFP-LC3-marked AVs showed a positive signal for mitochondria-derived NBD-PE signal in PS19 axons in the presence of glutamine (Figure 3G and H). PS19 axons without glutamine treatment consistently displayed lower AV density and, more importantly, most of these AVs were absent of NBD-PE made by mitochondria, further indicating that mitochondrial PE biosynthesis is not involved in autophagy in PS19 neurons under basal conditions. Moreover, by conducting time-lapse imaging in live PS19 axons under glutamine metabolism, we provided direct evidence showing dynamic AV formation from bioenergetically enhanced mitochondria in a 15-second recording period (Figure 3I). Together, these results suggest that glutamine-enhanced OXPHOS promotes AV biogenesis by powering mitochondrial PE biosynthesis in tauopathy neurons.

### Early stimulation of anaplerotic metabolism elevates OXPHOS activity and restores impaired PE biosynthesis in the mitochondria of tauopathy mouse brains

We then sought to address whether early stimulation of anaplerotic metabolism boosts autophagy activity for tau clearance in tauopathy mouse brains. Previous studies have shown that dietary supplementation of 4% glutamine induces neuroprotective effects on mice with no toxicity [58,67,68]. Our recent work revealed that OXPHOS deficiency in tauopathy mouse brains can be detected as early as 3-4 months of age [46]. Therefore, we supplemented drinking water with 4% glutamine for non-Tg and PS19 mice at 3 months of age. We first determined whether defective bioenergetics can be reversed in the brains of tauopathy mice supplemented with glutamine. We measured the OCR of mitochondria freshly isolated from the cortices of these mouse brains after 12 weeks of glutamine supplementation and found that mitochondrial bioenergetic activity was markedly reduced in PS19 mouse brains, relative to that in non-Tg mouse brains (Basal: p = 0.0027; Maximal: p = 0.0037; ATP-linked: p = 0.0015) (Figure 4A). Excitingly, we also detected significant increases in mitochondrial basal, maximal, and ATP production-linked respiration rates in glutamine-treated PS19 mice as compared to those of control PS19 mice, which were fed with regular water and exhibited OXPHOS deficits (Figure 4A). Thus, this result suggests that glutamine supplementation reverses impaired mitochondrial bioenergetics in PS19 mouse brains, supporting our reasoning to further investigate its impact on mitochondrial PE biosynthesis. We carried out lipidomics analysis of mitochondrial lipids extracted from the cortices of non-Tg and PS19 mouse brains with and without 12-week glutamine administration. The quantification of this data revealed marked decreases in the fold change of total mitochondrial PE species and the relative abundance of individual PE species in the PE class of mitochondria in PS19 mouse brains, compared to those of non-Tg mouse brains (p < 0.0001) (Figure 4B-D). Together with the data from OCR measurement (Figure 4A) and our imaging studies in live tauopathy neurons (Figure 3E and F), these findings consistently indicate impaired PE biosynthesis in the mitochondria of tauopathy mouse brains associated with OXPHOS deficits. In contrast, mitochondria in the brains of PS19 mice with glutamine supplementation displayed significant increases in the fold change of total PE species and the relative abundance of individual PE species (Figure 4B-D). Similar effects can be detected in the brain mitochondria of non-Tg mice under glutamine metabolism. Notably, the levels of PS, the PSD substrate [34], were also elevated in the brain mitochondria of glutamine-treated non-Tg and PS19 mice (p < 0.0001) (Figure S4A-C). These observations collectively support a critical role of OXPHOS activity in the control of mitochondrial PE metabolism and further indicate that stimulation of OXPHOS is the key to the rescue of tauopathy-linked PE biosynthesis defects in mitochondria.

**Figure 4.**
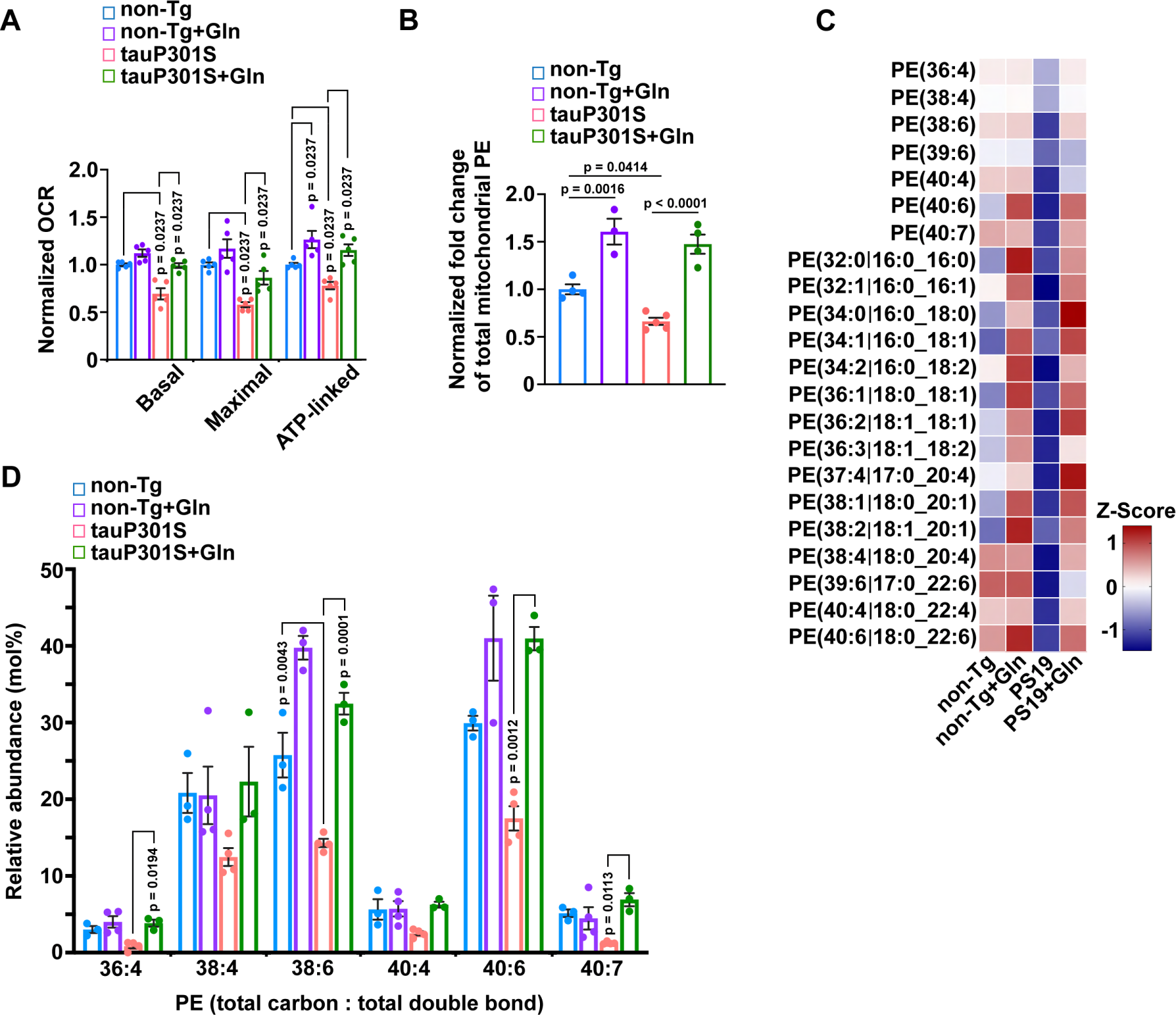
Early stimulation of anaplerotic metabolism elevates OXPHOS activity and increases PE biosynthesis in tauopathy mouse brain mitochondria. (A) Seahorse mitochondrial stress assay for OCR measurements of freshly isolated cortical mitochondria from the brains of non-Tg and PS19 mice at 6 months of age with and without 12-week glutamine supplementation. Basal, maximal, and ATP production-linked respiration rates of mitochondria purified from glutamine-administered non-Tg and PS19 mouse brains were normalized to those from control mice fed with glutamine-free water, respectively. OCR, oxygen consumption rate. (B) Lipidomics analysis of the fold change of total mitochondrial PE species in the brains of 6 months of age non-Tg and PS19 mice administered with water or 12-week glutamine. The changes of mitochondrial PE in the brains of PS19 control mice and non-Tg and PS19 mice supplemented with glutamine were normalized to that of non-Tg control mouse brains fed with water, respectively. (C) Heat map analysis for molecular phospholipid species of PE extracted from 6-month-old non-Tg and PS19 mouse brain mitochondria with and without glutamine administration. Rows represent the mean values for PEs and columns represent those of non-Tg or PS19 mice. (D) Relative abundance (mol%) of individual PE species in the total PE class of mitochondria in the brains of non-Tg and PS19 mice administered with water or glutamine. Data were quantified from five animals (A) and three to four animals (B-D) in each group, respectively. Data were expressed as the mean ± SEM and analyzed by Kruskal-Wallis test with Bonferroni’s correction (A) or one-way ANOVA with Bonferroni’s correction (B and D).

### Anaplerotic stimulation of OXPHOS enhances autophagy and reduces tau accumulation in tauopathy mouse brains

Given increased bioenergetics along with elevated PE in the brain mitochondria of glutamine-supplemented tauopathy mice (Figure 4), we performed three lines of experiments to examine whether enhanced mitochondrial PE biosynthesis boosts autophagy activity for tau clearance in these tauopathy mouse brains. First, our immunohistochemistry studies showed that LC3 appeared mostly in a diffuse pattern in the non-lipidated LC3-I form, which was present in the cytoplasm of the hippocampus of control PS19 mouse brains fed with glutamine-free water. We only detected few LC3-II-decorated AVs, shown as vesicular LC3-positive clusters (Figure 5A and B). Interestingly, the brains of PS19 mice administered with glutamine exhibited a marked increase in AV clusters in hippocampal CA3 and mossy fibers enriched with axons and presynaptic terminals. Our *in vitro* studies demonstrated that, in PS19 axons with high OXPHOS activity, newly formed AVs underwent predominantly retrograde transport toward the soma for lysosomal degradation (Figure 2G and H). We thus examined autolysosomes in the soma of hippocampal neurons in PS19 mouse brains with and without glutamine treatment. Glutamine administration significantly elevated the density of autolysosomes co-labeled with LC3 and LAMP1 antibodies and LC3-positive AVs or LAMP1-positive lysosomes relative to that of control PS19 mouse brains fed with regular water (Figure S5A and B). Second, we conducted transmission electron microscopy (TEM) analysis and examined presynaptic autophagy in these PS19 mouse brains at the ultrastructural level. Strikingly, presynaptic terminals in the brains of PS19 mice receiving glutamine treatment demonstrated initial AV (AVi)-like double-membrane structures near or directly derived from mitochondria, termed mitochondria-derived AVs (MDAVs), which were not readily observed in those of control PS19 mice without glutamine supplementation (Figure 5C and D). Besides increased presynaptic AV density, we also observed elevated distribution of mitochondria at presynaptic terminals in these glutamine-treated PS19 mouse brains (Figure 5C and D), which is consistent with our light imaging data obtained from cultured PS19 neurons (Figure 1C and D). Third, we biochemically assessed tau levels and autophagy activity in these mouse brains. We first examined different tau forms in detergent soluble protein lysates by running native gels and using Tau5 antibody against total tau. Relative to control PS19 mice fed with regular water, the brains of PS19 mice administered with glutamine demonstrated decreased tau levels (Figure 5E and F). Importantly, oligomeric tau, a toxic form of tau species, showed a greater reduction than monomeric tau, which is consistent with our *in vitro* data showing that OXPHOS-induced autophagy favors clearance of tau oligomers in tauopathy neurons (Figure 2C and D). Notably, in non-Tg mouse brains, glutamine supplementation did not lead to a significant change in tau levels and there were no detectable tau oligomers (Figure 5E and F). We thus performed a comprehensive analysis of control and glutamine-treated PS19 mouse brains for autophagy and tau proteins using phospho-tau antibodies (AT8, S202/T205, and PHF1, S396/S404) as well as antibodies against total Tau (Tau5) and unphosphorylated normal tau at the S202/T205 site (Tau1). OXPHOS stimulation in PS19 mouse brains led to increased LC3-II and reduced p62 (Figure 5G and H), which is in agreement with our imaging data (Figure 5A and B) and indicates enhanced autophagy activity. Furthermore, glutamine-treated PS19 mouse brains exhibited significant reductions in the levels of phospho-tau, recognized by both PHF1 and AT8 antibodies (Figure 5G and H). The data is in accord with the evidence of decreased tau oligomers revealed by native gels (Figure 5E and F). In addition, the levels of phospho-tau for both murine Tau (mTau) and human (hTau) showed reductions in these PS19 mouse brains (Figure 5I). As for normal tau, we observed an increase in mTau and no significant change in hTau. Thus, OXPHOS-stimulated reduction in total tau was mostly attributed to phospho-tau decrease (Figure 5G-I). These *in vivo* observations are consistent with our *in vitro* imaging and biochemical results (Figure 1C-F), collectively suggesting that OXPHOS-enhanced autophagy is selective for tau clearance and has protective effects at tauopathy synapses. To rule out model-specific effects, we included a second tauopathy mouse model—rTg4510. This well-characterized regulatable tauopathy model overexpresses 13 units of human mutant (P301L) tau downstream of a tetracycline-operon-responsive element (TRE) [5,69]. We supplemented drinking water for rTg4510 mice with 4% glutamine for 16 weeks starting at the age of 2 months. Consistent with the findings in glutamine-treated PS19 mouse brains, we observed similar increases in AVs and autolysosomes co-labeled by LC3 and LAMP1 antibodies in the neuronal soma of hippocampal CA1 regions of glutamine-treated rTg4510 mouse brains (Figure S5C and D). Together, these results suggest that OXPHOS stimulation-enhanced autophagy prevents pathological tau accumulation in tauopathy mouse brains.

**Figure 5.**
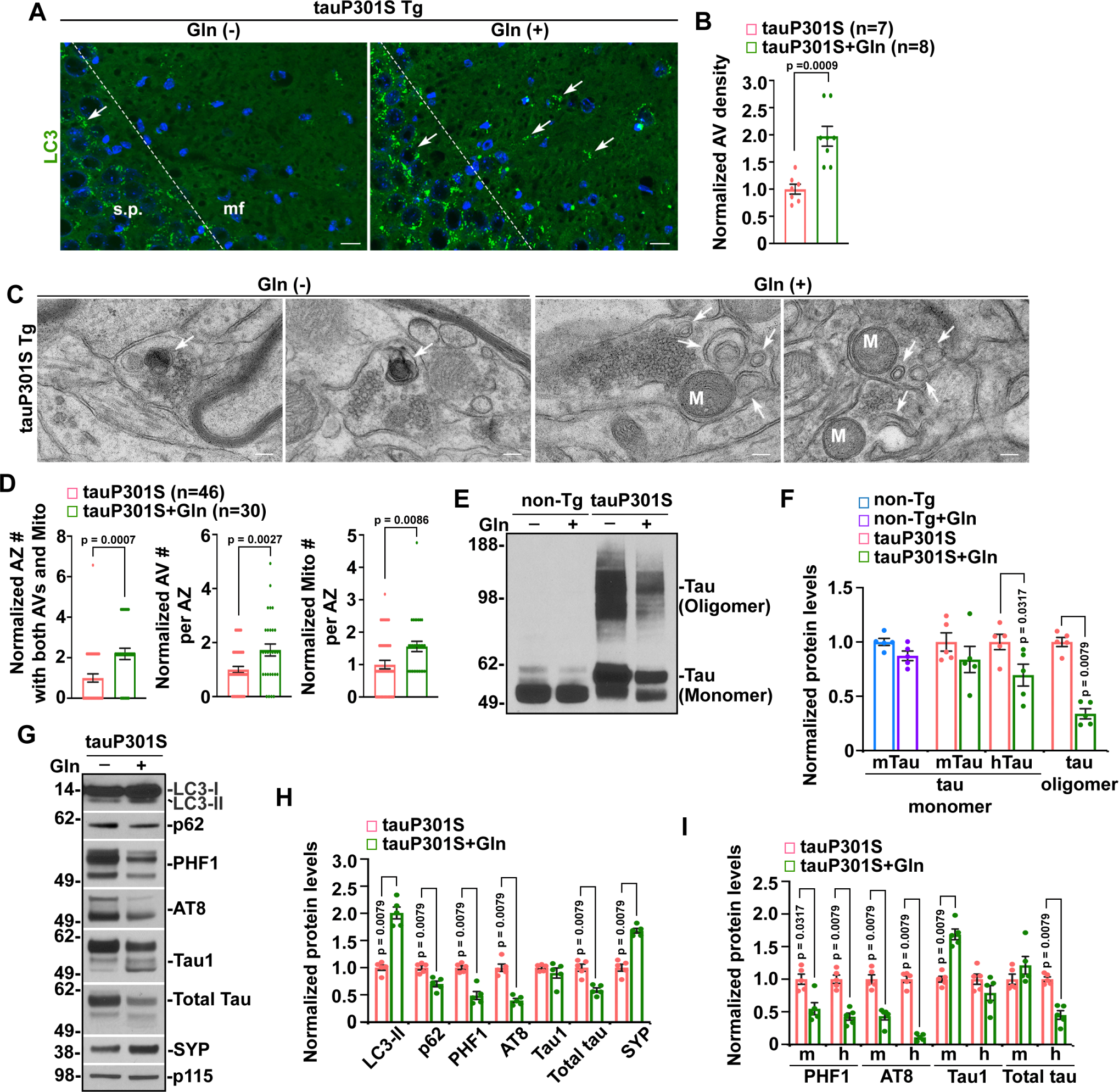
Autophagy enhancement coupled with tau reduction in metabolically reprogrammed tauopathy mouse brains. (A-B) Representative images (A) and quantitative analysis (B) of AV clusters (arrows) in the hippocampal regions of 9-month-old PS19 mouse brains with and without glutamine supplementation. Data were quantified and normalized to those of control PS19 mouse brains fed with glutamine-free water from a total number of imaging slice sections as indicated in parentheses (B) from two pairs of PS19 mice. s.p.: Stratum pyramidale; mf: mossy fibers. (C-D) Representative transmission electron microscopy (TEM) images (C) and quantitative analysis (D) of double-membrane initial AV (AVi)-like structures on or near mitochondria at the presynaptic terminals of control and glutamine-treated 6-month-old PS19 mouse brains. M: mitochondrion; arrows: AVs. Data were quantified from the total number of presynaptic terminals (n) as indicated in parentheses (D) from two pairs of PS19 mice. (E-F) Representative blots (E) and quantitative analysis (F) of monomeric and oligomeric tau levels revealed by a native gel in 9-month-old non-Tg and PS19 mouse brains with and without glutamine treatment that were normalized to those of control non-Tg littermate or PS19 mouse brains in the absence of glutamine. Data were quantified from five independent repeats with five mice in each group. mTau: murine Tau; hTau: human Tau. (G-I) Represent blots (G) and quantitative analysis (H and I) of autophagy and tau protein levels in detergent-extracted mouse brain lysates from 9-month-old PS19 mice administered with regular water or glutamine. The protein levels in the brains of PS19 mice with glutamine supplementation were normalized to p115 and to those in control PS19 mice fed with water, respectively (H and I). Data were quantified from five pairs of PS19 mice. m: murine; h: human. Data were expressed as the mean ± SEM and analyzed by Mann-Whitney U test (B, F, H, and I) or two-sided unpaired Student’s *t*-test (D). Scale bars: 10 μm (A) and 200 nm (C).

### Mitochondrial bioenergetic enhancement mitigates tau pathology, synapse loss, and neuronal death in tauopathy mouse brains

We then carried out more immunohistochemical staining to examine tau pathology in these bioenergetically enhanced tauopathy mouse brains. Stimulation of OXPHOS activity led to a robust reduction in phospho-tau and NFT-like pathologies revealed by AT8 and PHF1 antibodies in the hippocampus of PS19 mouse brains (Figure 6A and B). Next, we determined whether metabolic reprogramming offers neuroprotective effects in tauopathy mouse brains. Compared to non-Tg littermate mice, we found that both the area and the mean intensity of SYP-labeled presynaptic terminals were significantly reduced in the hippocampal mossy fiber regions of the brains of PS19 mice at the age of 9 months (SYP area: p < 0.0001; SYP mean intensity: p < 0.0001) (Figure 6C and D), which is in accord with previous studies from us and others [6,46,70,71]. Such a defect was remarkably alleviated in the brains of PS19 mice administered with glutamine (Figures 6C and D, 5G and H). It was worth noting that rTg4510 mice receiving glutamine supplementation also exhibited similar rescue effects on synapse loss in the hippocampal mossy fibers (SYP area: p < 0.0001; SYP mean intensity: p < 0.0001) (Figure S6A and B) as well as on tau pathology in the hippocampus (Figure S6C and D). Furthermore, we examined whether stimulation of mitochondrial bioenergetics results in neuroprotective effects in tauopathy mouse brains. PS19 mice exhibited a significant decrease in neuronal density in the hippocampal CA3 regions, relative to non-Tg littermate controls. Glutamine treatment-enhanced bioenergetic activity mitigated neuronal death, as evidenced by the increased number of NeuN-positive neurons in this region of PS19 mouse brains (p < 0.0001) (Figure 6E and F). Thus, the *in vivo* evidence from tauopathy mouse brains indicates the protective effects of early metabolic stimulation against tauopathy-associated tau pathology and neurodegeneration.

**Figure 6.**
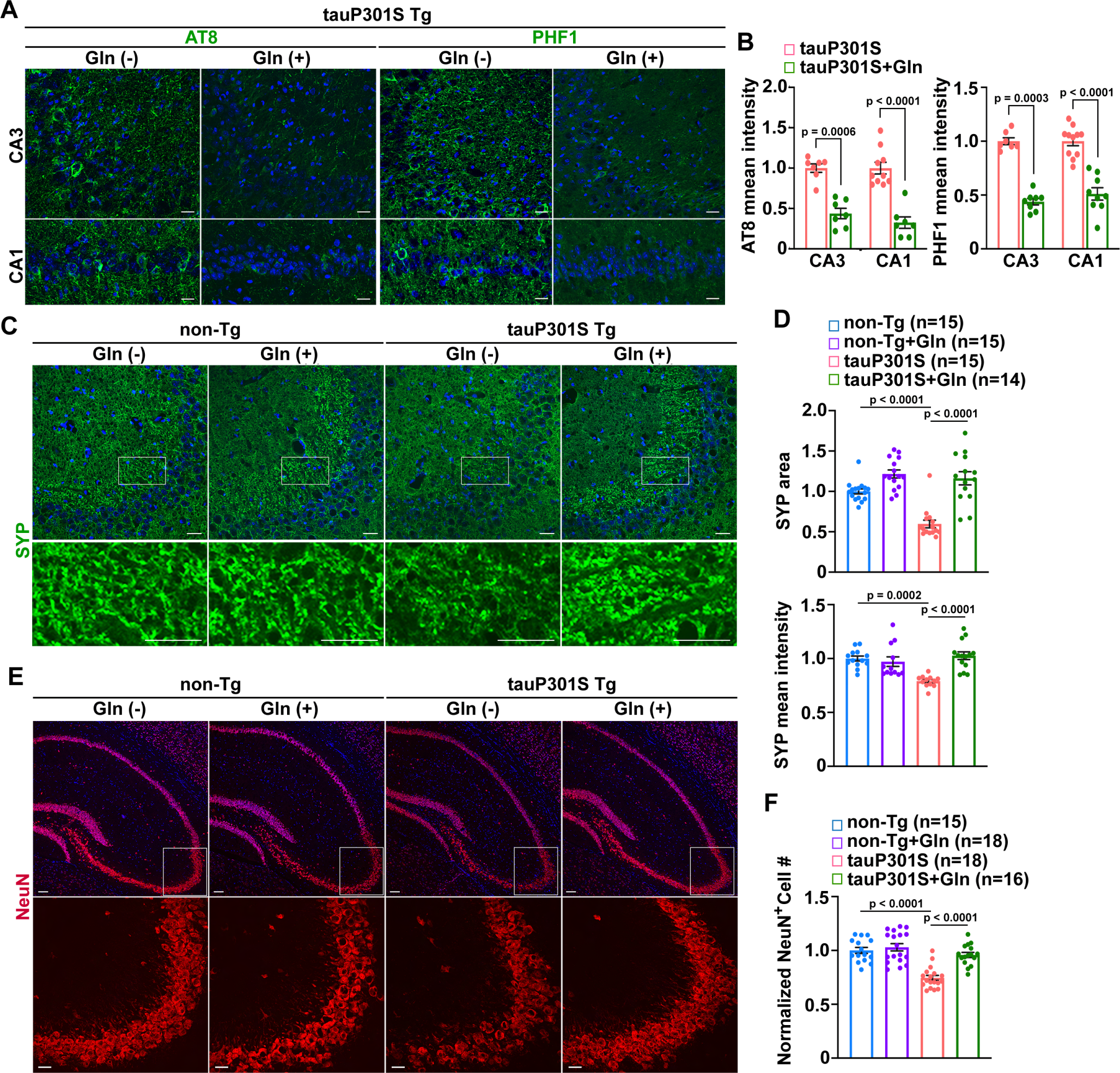
OXPHOS stimulation attenuates tau pathology and alleviates synapse loss and neurodegeneration in tauopathy mouse brains. (A-B) Representative images (A) and quantitative analysis (B) of tau accumulation/aggregation in the hippocampus of 9-month-old PS19 mouse brains with and without glutamine supplementation (n = 7-11 mouse brain slice sections in each group from two pairs of PS19 mice). The mean intensities of AT8 and PHF1 antibody-marked phospho-tau in the hippocampal CA3 and CA1 areas were quantified and normalized to those of control PS19 mouse brains fed with regular water, respectively. (C-D) Representative images (C) and quantitative analysis (D) of presynaptic terminal densities in the hippocampal mossy fibers of 9-month-old non-Tg and PS19 mice administered with water or glutamine. The area and the mean intensity of SYP-marked presynaptic terminals were quantified and normalized to non-Tg control mice with no glutamine treatment, respectively. (E-F) Representative images (E) and quantitative analysis (F) of neuron density in the hippocampal regions of non-Tg and PS19 mouse brains with and without glutamine administration. The number of NeuN-labeled neurons in hippocampal CA3 areas marked by rectangles was quantified and normalized to that of non-Tg mice in the absence of glutamine. Data were quantified from a total number of imaging slice sections as indicated in parentheses (D and F) from three mice per group. Data were expressed as the mean ± SEM and analyzed by Mann-Whitney U test (B) or one-way ANOVA with Bonferroni’s correction (D and F). Scale bars: 10 μm (A), 25 μm (C and the bottom panel in E), and 250 μm (the top panel in E).

### Metabolic reprogramming leads to improvement of behavioral performance in tauopathy mice

Given OXPHOS enhancement-induced neuroprotective effects in tauopathy mouse brains (Figures 6 and S6), we postulated that glutamine metabolism ameliorates learning and memory deficits in tauopathy mice. We asked whether early metabolic stimulation prevents these behavioral abnormalities, which are readily detectable in PS19 mice. In the open field test, 8-month-old PS19 mice displayed greater locomotion and less anxiety-like behavior, as evidenced by increased travel distance in the open field, and more time spent in the field center, an indication of reduced anxiety. This suggests that tauopathy mice were more hyperactive (total movement: p = 0.0205; center time: p = 0.0414) (Figure S7), a finding that is consistent with previous studies [72-74]. Glutamine supplementation showed no significant effect on the hyperactivity of these PS19 mice (p > 0.05) (Figure S7). Phospho-tau accumulation in the prefrontal cortex and amygdala was proposed to induce hyperactive behavior [6,75,76]. Thus, our results suggest the persistence of brain regions driving hyperlocomotion behavior, which is resistant to bioenergetic enhancement.

We next performed the three-chamber test to assess the impact of glutamine treatment on the sociability and social recognition memory [72,77-79]. Animals with normal sociability show a preference for the chamber with a stranger mouse over an empty chamber, a phenomenon indicated by the longer time they would spend in the stranger chamber. PS19 mice showed comparable social interaction relative to non-Tg littermate controls (Figure 7A). During the social novelty preference/recognition test that requires normal hippocampal function, non-Tg mice spent significantly longer time exploring the chamber containing a novel, stranger mouse relative to that containing a familiar one. However, PS19 mice failed to distinguish the stranger mouse from the familiar mouse and spent comparable time with the stranger mouse relative to the familiar mouse (p = 0.3823) (Figure 7A). This observation is consistent with previous studies [72], suggesting that PS19 mice have impaired social memory. Importantly, this phenotype was significantly reversed in glutamine-treated PS19 mice, as evidenced by more time these animals spent with the stranger relative to the familiar mouse (p = 0.0111). Therefore, these results demonstrate the rescue effects of early metabolic stimulation on tauopathy-associated social recognition memory defects.

**Figure 7.**
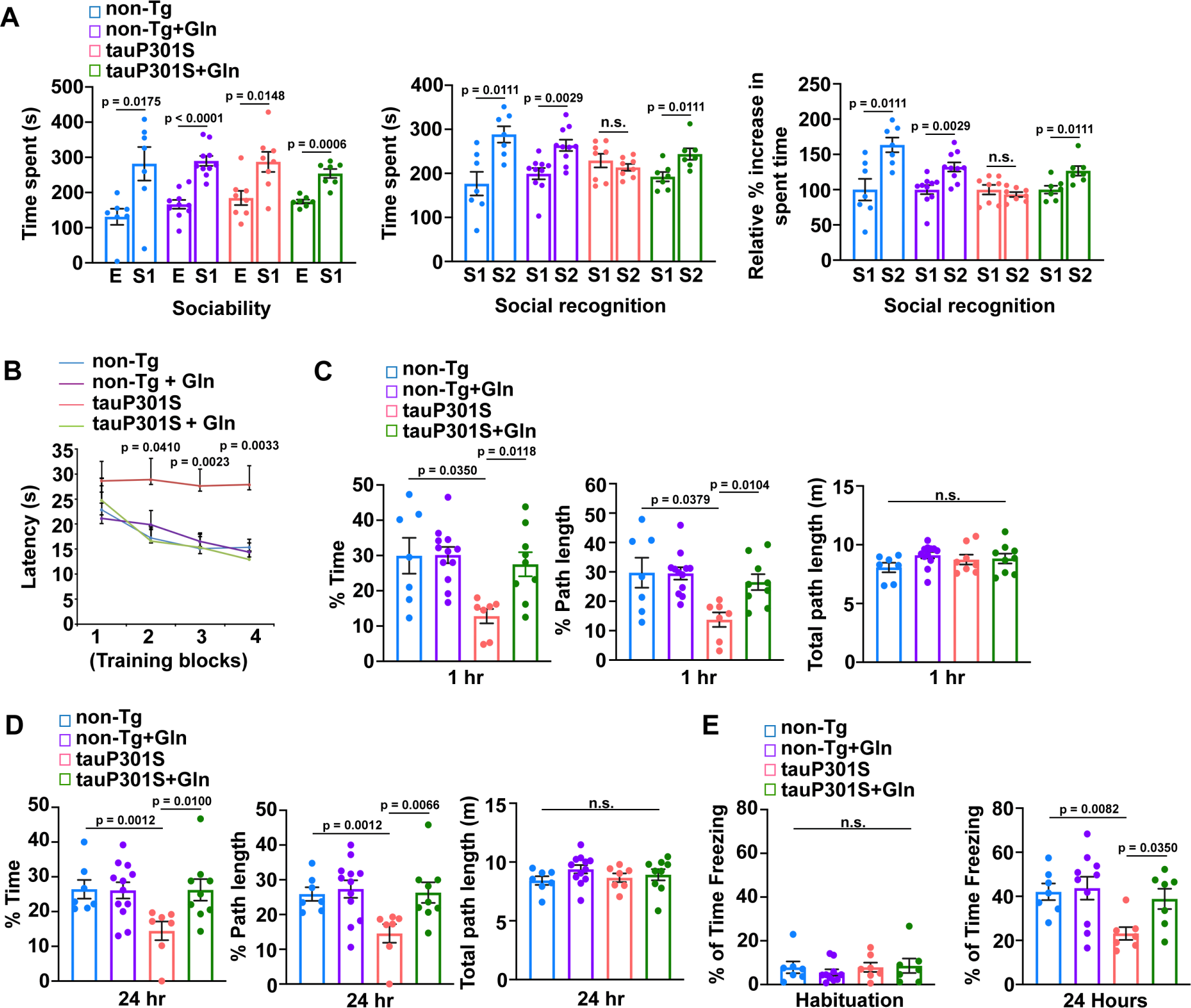
Metabolic reprogramming leads to improvement of behavioral performance in tauopathy mice. (A) Sociability and social recognition memory assay performed in 8-month-old non-Tg and PS19 male mice administered with regular glutamine-free water or glutamine (n = 7-11 male mice per group). n.s.: non-significant. (B-D) Morris water maze test of 8-month-old PS19 male mice and non-Tg littermates with and without glutamine administration (n = 7-12 male mice per group). (E) Contextual fear conditioning task performed in 8-month-old non-Tg and PS19 male mice supplemented with regular water or glutamine (n = 7-10 male mice per group). Data were shown as the mean ± SEM and analyzed by Mann-Whitney U test (A) or Kruskal-Wallis one-way analysis of variance test with Bonferroni’s correction (B-E).

To examine the impact of metabolic reprogramming on spatial learning and memory, we conducted the Morris water maze (MWM) test in 8-month-old PS19 mice and non-Tg littermates with and without glutamine administration. Mice were trained to find a hidden platform across four training blocks (Blocks 1-4) on a single day and with each block consisting of 3 to 4 acquisition trials, followed by memory testing in a 60-second probe trial at 1 hour or 24 hours after the last hidden-platform training, respectively [80,81]. Non-Tg mice receiving water supplemented with glutamine or those receiving glutamine-free water displayed similar latencies to find the platform during the hidden-platform training phase and favored the target quadrant in the probe trials (non-Tg vs. non-Tg + Gln) (Figure 7B-D), indicating that glutamine metabolism causes no adverse effects in learning and memory in non-Tg mice. In contrast, PS19 mice fed with water alone performed poorly in both the acquisition phase and the memory probe tests, as reflected by longer latencies on training blocks 2-4 (block 2: p = 0.0258; block 3: p = 0.0010; block 4: p = 0.0091) and shorter path lengths and less time in the target quadrant during probe trials that were conducted at 1 hour or 24 hours after training trials (non-Tg vs. PS19). However, stimulation of OXPHOS significantly improved learning and enhanced memory retention in PS19 mice to levels that showed significant difference from those of control PS19 mice (block 2: p = 0.0137; block 3: p = 0.0020; block 4: p = 0.0012) (PS19 vs. PS19 with glutamine) (Figure 7B-D), suggesting that metabolic reprogramming in PS19 mice significantly ameliorates deficits in spatial learning and memory tests.

Using a contextual fear conditioning task [82], we further examined the effects of glutamine treatment on contextual memory in mice at 8 months of age. PS19 mice with or without glutamine treatment showed no significant difference in freezing from non-Tg littermates on the training task during habituation to the apparatus prior to training with foot shock exposure (p = 0.6980), but showed a significant difference in freezing 24 hours after the training (p = 0.0195) (Figure 7E). Subsequent to training, non-Tg mice administered with and those without glutamine supplementation exhibited contextual fear learning, with 42.05% and 43.67% of the time spent “freezing” in the operant chamber in anticipation of the shock, respectively (Figure 7E). However, PS19 mice showed significant defects in this task, spending only 23.14% of the time freezing (p = 0.0082). These results are consistent with previous studies [71,72] and suggest defects in hippocampus-dependent memory in this tauopathy mouse model. Importantly, PS19 mice under glutamine supplementation exhibited significant amelioration of the learning deficit (p = 0.0350) compared to PS19 mice without glutamine treatment. This result is in line with OXPHOS enhancement-induced rescue effects on synaptic defects and neuronal death, indicating protective effects against hippocampus-dependent memory deficits in PS19 mice.

## Discussion

Mitochondrial bioenergetic failure and energy deficits are early and prominent features in tauopathy brains [46-49]. Autophagy defects have also been implicated in patient brains and cellular models of tauopathy [22-25]. Given that autophagy is an ATP-dependent process [83,84], a fundamental question exists as to whether defective mitochondrial energy metabolism plays a role in the pathogenesis of tauopathy-linked autophagy dysfunction. In addressing this question, the findings from the current study support the mechanistic link between OXPHOS deficiency and autophagy failure in tauopathy and further demonstrate that enhancement of mitochondrial bioenergetics is key to the rescue of autophagy defects and tauopathy-related pathologies.

We and others have established that axons and synaptic terminals are the primary sites for AV formation, which facilitates the sequestration of autophagic cargos/substrates within AVs. These nascent AVs return the engulfed cargos through retrograde transport for lysosomal clearance in the soma. Such a mechanism is critical for the prevention of toxic accumulation at synaptic terminals [15-21]. Pathological tau is targeted by autophagy for clearance [7-10]. Previous findings from us and others highlighted that phospho-tau is highly concentrated at synaptic terminals in tauopathy models [5,46,85-90], suggestive of autophagy failure at tauopathy synapses. Our recent studies demonstrated that synaptic mitochondrial deficits occur early in tauopathy [46]. These findings indicate a possible link between mitochondrial bioenergetic deficiency and defects in the autophagy-mediated removal of phospho-tau from tauopathy synapses. The present study has shown that stimulation of anaplerotic metabolism boosts OXPHOS activity in tauopathy neurons and has further provided multiple lines of evidence showing the connection between OXPHOS enhancement and increased AV biogenesis.

First, we have revealed that OXPHOS stimulation leads to an increased number of AVs in distal PS19 axons (Figure 2E and F). Moreover, our time-lapse live cell imaging data directly demonstrated that AVs are formed from a mitochondrion in a bioenergetically enhanced PS19 axon (Figure 3I). These *in vitro* data are consistent with results from our TEM analysis showing drastic AV formation on or near mitochondria at the presynaptic terminals of PS19 mouse brains supplemented with glutamine (Figure 5C and D). Second, we have shown that nascent AVs in glutamine-treated PS19 axons undergo retrograde movement toward the soma for lysosomal degradation (Figure 2G and H). In line with this data, we found a significant increase in AVs in the soma of PS19 neurons treated with glutamine (Figure S2C and D). In addition, previous studies have established that blocking lysosomal proteolysis activity causes autophagosomes to lose dynein and stall in the axons [15,91]. Indeed, we observed an augmentation of AV increase in glutamine-treated PS19 axons upon lysosomal inhibition (Figure 2E and F), which consistently suggests OXPHOS-enhanced AV biogenesis. Third, we have shown that elevated AV biogenesis in glutamine-treated PS19 axons is abrogated upon inhibition of mitochondrial PE biosynthesis (Figures 3C and D, S3E and F). These observations suggest that OXPHOS-enhanced AV biogenesis in PS19 axons rely on the mitochondrial PE pool. Fourth, glutamine-enhanced OXPHOS is found to result in increased AVs in the axons of non-Tg neurons (Figure S2E-H). Taken together, our *in vitro* and *in vivo* findings in normal and tauopathy neurons consistently suggest the link between mitochondrial bioenergetic enhancement and increased AV biogenesis.

Notably, PS19 mouse brains supplemented with glutamine exhibited a selective reduction in pathological tau but not normal tau (Figure 5G-I), indicating that high OXPHOS activity-induced autophagy favors pathological tau clearance. This is consistent with our *in vitro* and *in vivo* imaging and biochemical results demonstrating that glutamine-treated PS19 neurons showed a decrease in phospho-tau but not mitochondrial and neuronal protein levels (Figure 1C-F). Moreover, autophagy is a lysosome-dependent pathway[11,26]. Despite controversial results from earlier studies in isolated rat hepatocytes regarding the impact of intracellular concentration of ATP on lysosomal activity [83,84], we have observed an increase in LAMP1 positive-lysosomes/autolysosomes in the soma of hippocampal neurons of PS19 mice administered with glutamine. Thus, it is likely that bioenergetic stimulation-elevated cellular ATP levels also promote lysosomal proteolytic activity, thereby contributing to pronounced tau clearance by autophagy in tauopathy neurons. Combined, these results support the view that mitochondrial energy metabolism is critical for autophagy-mediated clearance of pathological tau, highlighting a pivotal role of bioenergetic defects in toxic tau buildup in tauopathy.

Many studies have highlighted the critical role of pathological tau in perturbing synaptic function, which triggers cognitive and behavioral decline in AD and other tauopathies [5,85-90]. Our results have demonstrated that bioenergetic enhancement-mediated rescue effects on autophagy dysfunction leads to an attenuation of tau pathology, thereby counteracting synaptic damage in tauopathy mouse brains. Neurodegeneration is thought to begin with the loss of presynaptic terminals and proceed retrogradely in a dying-back process. Indeed, we have further shown that early stimulation of mitochondrial bioenergetics protects against neurodegeneration and memory deficits in tauopathy mice. It should be noted that in three separate tests of cognitive behavior−social recognition, spatial navigation, and contextual fear learning−the results are consistent in demonstrating that OXPHOS enhancement rescues cognitive deficits in tauopathy mice. Given that various mechanisms have been proposed regarding the pathological role of mitochondrial defects in the pathogenesis of tauopathy [39,41,42,92], the current results support a new premise that mitochondrial bioenergetic deficits play a crucial role in tauopathy-linked autophagy failure and the buildup of toxic tau, contributing to synaptic deterioration and cognitive impairment.

Most studies to date investigating neuronal autophagy have focused on the mechanisms underlying compartmentalized AV formation and maturation, the molecular machinery that is involved, and the process of autophagic cargo selection and elimination [15,17,18,93-97]. Given that the origin of AV and the membrane sources for AV biogenesis have only been under extensive investigation in recent years [11,26,98], how such mechanisms participate in neuronal autophagy regulation remains poorly understood. Studies have shown that PE is a main constituent of the AV membrane and the supply of PE is a limiting factor for autophagy activity [30]. Therefore, reductions in cellular PE compromise AV formation and consequently restrict the availability of AVs for autophagic cargos which need to be sequestered within AVs to allow autophagy progression [30,99]. Previous work implies that PE is mainly synthesized in both the ER and mitochondria [34-36]. Interestingly, pools of PE from the ER and mitochondria are compartmentalized and do not mix freely within cells [100-102]. Several studies have demonstrated that PE biosynthesis either in the ER or in mitochondria can provide PE for AV biogenesis in non-neuronal cells under starvation conditions [30,63]. However, studies in DRG neurons reported that the ER was the primary source of membrane supply for AV formation [16], raising a question as to whether PE made in the ER might mediate OXPHOS-induced autophagy in tauopathy neurons. Unexpectedly, the current study has shown that inhibition of the CDP-ETN pathway-associated PE biosynthesis in the ER fails to block autophagy stimulated by OXPHOS. We have further provided *in vitro* and *in vivo* evidence demonstrating that OXPHOS-induced autophagy in tauopathy neurons rely on PE synthesized in mitochondria through the PSD pathway. Together, in addition to the role of the ER in autophagy in DRG neurons [16], our findings uncover that mitochondria are also critically involved in AV biogenesis for neuronal autophagy. It is important to note that, in normal neurons, anaplerotic stimulation of OXPHOS also led to a robust increase in autophagy activity by promoting AV biogenesis (Figure S2E-H). Thus, the current study provides new insights into a crucial role of mitochondrial bioenergetics-powered PE biosynthesis in the regulation of autophagy activity for cargo clearance in both normal and disease neurons.

It is noteworthy that ER-supported autophagy is functional and unaffected by mitochondrial bioenergetic status in tauopathy neurons. Enhancement of ER PE biosynthesis by incubation of tauopathy neurons with ETN, the substrate of the CDP-ETN pathway in the ER [34-36], effectively increased the amplitude of AV biogenesis in the axons in the absence of glutamine. Moreover, we did not observe an additional increase or reduction in autophagic flux in OXPHOS-enhanced tauopathy neurons in the presence of ETN or meclizine which inhibits ER PE biosynthesis [62] (Figure S3C and D). Our data are in accord with previous studies showing a key role of the ER in autophagy in DRG neurons under basal conditions [16], and further indicate that basal autophagy mediated by ER-dependent PE biosynthesis is less likely subject to the changes in mitochondrial bioenergetic status in tauopathy neurons. More importantly, ER PE pool-supported autophagy activity is much less potent than autophagy dependent on the mitochondrial PE pool and is insufficient in curbing toxic tau accumulation in tauopathy neurons. These observations collectively suggest that PE biosynthesis in mitochondria, but not in the ER, relies on OXPHOS activity and further support the notion that autophagy deficits and abnormal pathological tau accumulation in tauopathy neurons are largely attributed to defects in OXPHOS-powered PE biosynthesis in mitochondria as a consequence of mitochondrial bioenergetic deficiency.

Several studies demonstrated the ATP-dependent nature of PE metabolism/biosynthesis by depleting cellular ATP or adding exogenous ATP in permeabilized cell lines [38,66]. However, direct evidence demonstrating the energy source that powers ATP-dependent PE biosynthesis remains lacking. In the current study, we addressed this fundamental question by applying more physiological conditions such as cultured primary disease neurons and established tauopathy mouse models. We have demonstrated that, in tauopathy neurons, ATP production through OXPHOS fuels PE biosynthesis in mitochondria, but not in the ER. Results from our light imaging studies in live tauopathy axons and the lipidomics analysis of tauopathy mouse brain mitochondria consistently showed that mitochondrial PE biosynthesis became impaired when OXPHOS deficits emerged. Importantly, stimulation of anaplerotic metabolism in tauopathy neurons restored defective OXPHOS and led to a marked increase in the levels of mitochondrial PE, suggesting that OXPHOS activity powers PE metabolism/biosynthesis in mitochondria. It is also worth noting that tauopathy mouse brains with OXPHOS deficits exhibited a significant reduction in mitochondrial PS (Figure S4A-C), the specific substrate for PE biosynthesis in mitochondria [34]. Previous work has revealed that, while PS is synthesized in an ATP-independent fashion in the ER, ATP is required for the import of newly made PS into the IMM, the rate-limiting step for PSD-mediated conversion of PS to PE in mitochondria [38,103]. Our results support the view that OXPHOS-supplied ATP serves as the energy source fueling PS trafficking to the IMM. In line with this view, we observed an increase in mitochondrial NBD-PE fluorescence in OXPHOS-enhanced tauopathy neurons. Several studies have shown that, while the exogenous NBD-PS is efficiently partitioned into the outer mitochondrial membrane (OMM) in an ATP-independent fashion [66], the import of NBD-PS into the IMM where NBD-PS is converted to NBD-PE is ATP-dependent, which also allows the NBD fluorescent signals to be stabilized in mitochondria [63-66]. Otherwise, cytosolic NBD-PS stays unstable and is broken down by PS-specific phospholipase A1 and A2 [104-106]. Thus, in glutamine-treated tauopathy neurons, elevated NBD-PE fluorescence in mitochondria could be the result of enhancement of ATP-dependent transport of exogenous NBD-PS to mitochondria, leading to increased PE metabolism—PS conversion to PE in the IMM (Figure 3E and F). This *in vitro* data is also consistent with the evidence from lipidomics analysis showing elevated mitochondrial levels of both PS and PE in metabolically enhanced tauopathy mouse brains. Collectively, these findings imply that OXPHOS deficiency disrupts mitochondrial PE metabolism/biosynthesis by impairing ATP-dependent PS translocation to the IMM, leading to defects in AV biogenesis, and thus autophagy failure, in tauopathy neurons.

In summary, we delineate a new mechanism of mitochondria-mediated regulation of neuronal autophagy. Our work advances current knowledge by demonstrating an essential role of OXPHOS-powered mitochondrial PE metabolism/biosynthesis in neuronal autophagy under pathophysiological conditions and provides new mechanistic insights into the link between mitochondrial bioenergetic failure and autophagy deficits in tauopathy. In addition, the results have broader relevance to understanding of other neurodegenerative diseases and aging associated with mitochondrial defects, autophagy dysfunction, and toxic accumulation. Given that the efforts to develop Aβ-based therapies have largely failed [107,108], the present study may also benefit the development of new therapeutic strategies for treating AD and other tauopathy diseases.

## Methods

### Mouse line and animal care

TauP301L (rTg4510) and tauP301S (PS19) mouse lines [5,6] were purchased from the Jackson Laboratory. All animal procedures were carried out following NIH guidelines and were approved by the Rutgers Institutional Animal Care and Use Committee. The animal facilities at Rutgers University are fully Association for Assessment and Accreditation of Laboratory Animal Care accredited.

### Transfection of cultured cortical neurons

Mouse cortical neurons were prepared from cortical tissues dissected from E18-19 mouse embryos (sex: random) using the Papain method as previously described [20,109-113]. In brief, after dissociation by papain (Worthington), cortical neurons were resuspended in a plating medium (5% FBS, insulin, glutamate, G5 and 1 × B27) supplemented with 100 × L-glutamine in Neurobasal medium (Invitrogen) and plated at a density of 100,000 cells per cm^2^ on polyornithine- and fibronectin-coated coverslips (Carolina Cover Glasses) or 35 mm dishes. 24 hours after plating, neurons were maintained in conditioned medium with half-feed changes of neuronal feeding medium (1 × B27 in Neurobasal medium) every 3 days. Primary cortical neurons were cultured from breeding mice of PS19 line with non-Tg controls [6]. Genotyping assays were performed following culture plating to verify mouse genotypes. We examined both PS19 and non-Tg neurons derived from their littermates. Non-Tg and PS19 neurons were transfected with various constructs at DIV5-7 using Lipofectamine 2000 (Invitrogen) followed by time-lapse imaging 10-14 days after transfection prior to quantification analysis.

### Anaplerotic stimulation of OXPHOS in cultured cortical neurons

Cortical neurons at DIV10-14 were treated for 24 hours with 10 mM glutamine or 5 mM aspartic acid in Neurobasal-A medium (Invitrogen, Cat# A24775-01) supplemented with 1 mM pyruvate and 1 mM lactate. Control neurons were incubated in the medium in which not only 10 mM glutamine or 5 mM aspartic acid was replaced by 25 mM glucose, but also pyruvate and lactate were not supplemented. Treated neurons were subjected to imaging and biochemical analyses, respectively.

### Western blot

Harvested cortical neurons or mouse brain tissues were lysed with ice-cold lysis buffer [1% Triton X-100, 8 μg/ml aprotinin, 10 μg/ml leupeptin, 0.4 mM PMSF in TBS with 1 × PhosSTOP (Millipore/Sigma, Cat# 4906837001)]. Protein quantification of cell lysates or tissue homogenates was performed by BCA assay (Pierce Chemical Co.). Equal amounts of proteins were loaded and resolved by 4-12% Bis-Tris NuPAGE protein gel and analyzed by sequential western blots on the same membranes after stripping between each application of antibody. For semi-quantitative analysis, protein bands detected by ECL were scanned into Adobe Photoshop 2022, and analyzed using NIH ImageJ. Care was taken during exposure of the ECL film to ensure that intensity readouts were in a linear range of standard curve blot detected by the same antibody.

### Determination of cellular and mitochondrial ATP levels in live neurons

mitAT1.03 or AT1.03 was transfected into cortical neurons at DIV5-6 using Lipofectamine 2000. The neurons were imaged using an Olympus FV3000 microscope at DIV10-14. The FRET signal (YFP/CFP emission ratio) was measured and calculated from the individual cells expressing mitAT1.03 or AT1.03, which reflects the mitochondrial or cytoplasmic ATP levels.

### Assessment of mitochondrial phosphatidylethanolamine (PE) metabolism in live neurons

1-oleoyl-2-{6-[(7-nitro-2-1,3-benzoxadiazol-4-yl)amino]hexanoyl}-sn-glycero-3-phosphoserine (Avanti Polar Lipids, Inc, Cat# 810194C) was mixed with DOPC 18:1 (D9-Cis) PC 1,2-di-(9Z-octadecenoyl)-sn-glycero-3-phosphocholine at a 20:80 ratio (Avanti Polar Lipids, Inc, Cat# 850375). After removal of the chloroform in the mixture, the NBD-PS was reconstituted to a 2.2 mM stock in sterile PBS. NBD-PS at a 0.22 mM concentration was added to cultured neurons transfected with DsRed-Mito or mRFP-LC3 and incubated for 30 min at 37°C. Imaging was performed at time 0 or 2.5 hours to 3 hours post-NBD-PS loading.

### Live neuron imaging

During live-cell imaging, neurons were maintained in custom-ordered Hibernate medium (HE-CUSTOM no glucose, no sodium pyruvate, no phenol red) supplemented with 10 mM glutamine, 1 mM pyruvate, and 1 mM lactate following 24-hour glutamine incubation. As for control neurons, Hibernate medium supplemented with 10 mM glucose, 0.5 mM glutamine, 1 mM pyruvate, and 1 mM lactate was used. The temperature was maintained at 37°C with an air stream incubator. Cells were visualized with a 60x oil immersion lens (1.3 numerical aperture) on an Olympus FV3000 confocal microscope, using 458 nm excitation for CFP, 488 nm for GFP, YFP, or NBD, and 559 nm for DsRed or mRFP. We selected the axons with at least 150 μm in length and more than 200 μm away from the soma of non-Tg or PS19 neurons. Axonal processes were selected as we previously reported [21,46,109-111]. Briefly, axons in live images were distinguished from dendrites based on known morphologic characteristics: greater length, thin and uniform diameter, and sparse branching [114]. Only those that appeared to be single axons and separate from other processes in the field were chosen for recording axonal mitochondrial transport. Regions, where crossing or fasciculation occurred, were excluded from the analysis.

For time-lapse imaging, tine-lapse sequences of 1,024 × 1,024 pixels (8 bit) were collected at 3-sec intervals with 1% intensity of the laser to minimize laser-induced bleaching and cell damage while maximizing pinhole opening. Time-lapse images were captured at a total of 100 frames. Recordings were started 6 min after the coverslip was placed in the chamber. The stacks of representative images were imported into NIH ImageJ software and converted to kymographs. To trace the anterograde or retrograde movement of axonal AVs and to count stationary ones, kymographs were made as described previously [110,115] with extra plug-ins for NIH ImageJ. Briefly, we used the “Straighten” plugin to straighten curved axons and the “Grouped ZProjector” to z-axially project re-sliced time-lapse images. The height of the kymographs represents recording time (300 sec unless otherwise noted), while the width represents the length (μm) of the axon imaged. Counts were averaged from 100 frames for each time-lapse image to ensure the accuracy of stationary and motile events. Vertical lines represent stationary organelles, while slanted lines to the right (negative slope) represent anterograde movement, and to the left (positive slope) indicate retrograde movement. An AV was considered stationary if it remained immotile (displacement ≤ 5 μm). Relative motility of vesicles or organelles is described as the percentage of anterograde, retrograde, or stationary events of total vesicles or organelles.

### Glutamine supplementation in tauopathy mice

4% glutamine (Sigma, Cat# G5792) in sterile tap water was made fresh daily and offered as the sole source of drinking water for 5 consecutive days (Monday through Friday), as previously described [58,67,68]. Control mice were fed with only sterile tap water. To avoid undue stress from elevated ammonia concentrations, all mice drank glutamine-free water 2 days each week (Saturday and Sunday). Glutamine administration in PS19 mice started at 3 months of age and continued until the end of the studies with the animals at the age of 6 months for OCR measurements and lipidomics analysis of mitochondria and transmission electron microscopy (TEM) studies, 9 months for biochemical and histological analyses, and 8 months for behavioral assays, respectively. rTg4510 mice were supplemented with glutamine at 2 months of age and for 16 weeks prior to histological studies. While male mice only were assessed for cognitive and emotional behavior, both male and female mice were used for other experiments.

### Respiration measurement of mouse brain mitochondria

For oxygen consumption rate (OCR) measurement, cortical tissues from non-Tg or PS19 mice with and without glutamine supplementation were used as a source for mitochondrial preparations. The tissues were rinsed in ice-cold mitochondrial isolation buffer [70 mM sucrose, 210 mM mannitol, 5 mM HEPES, 1 mM EGTA, 0.5% (w/v) fatty acid free BSA, pH 7.2], then homogenized using a glass Dounce tissue grinder (10 strokes with loose pestle, 10 strokes with tight pestle). Mitochondria were then freshly isolated from these tissue homogenates using a differential centrifugation method [46,58,116]. Briefly, the homogenate was centrifuged at 1,000 × g for 10 min at 4°C. Following centrifugation, the supernatant was transferred to a separate tube and centrifuged at 8000 × g for 15 min at 4°C and washed once more with the same buffer and centrifugation procedure. Protein concentrations were measured using BCA protein assay. Seahorse XF24 Analyzer (Agilent Technologies, CA) was used to measure bioenergetic function in freshly isolated mitochondria from the cortex and hippocampus. The XF24 creates a transient 7-μl chamber in specialized 24-well microplates that allows for the OCR to be monitored in real time [116]. 40 μg of isolated mitochondria was added to each well in 50 μl of mitochondrial assay solution [MAS: 70 mM sucrose, 220 mM mannitol, 10 mM KH_2_PO_4_, 5 mM MgCl_2_, 5 mM HEPES, 1 mM EGTA, 0.2% (w/v) fatty acid free BSA, pH 7.2]. The plate was then transferred to a centrifuge equipped with a swinging bucket microplate adaptor and spun at 2,000 × g for 20 min at 4°C. After centrifugation, 450 μl of MAS with pyruvate (10 mM) and malate (2 mM) was added to each well. Oligomycin (1 μM), FCCP (4 μM), and rotenone (1 μM) + antimycin A (1 μM) were injected sequentially through ports in the Seahorse Flux Pak cartridges. Each loop began with mixing for 1 min, followed by OCR measurement for 3 min. This allowed determination of the basal level of oxygen consumption, maximal respiration, as well as ATP-linked respiration [46,58,116].

### Lipidomics analysis of mouse brain mitochondria

After isolation of mitochondria from cortical tissues in non-Tg or PS19 mice with and without glutamine administration, isolated mitochondria equivalent to 1 mg protein were mixed with 250 μl methanol before adding 5 mL 0.1M Hydrochloric acid (Sigma, Cat# 1090601000) and 100 µL SPLASH (Avanti Polar Lipids, Cat# 330707,) to make Extraction Solvent. 1 ml MTBE (Sigma, Cat# 34875,) was added to 0.25 mL Extraction Solvent, which resulted in two layers after vortexing for 30 sec. 600 µl from the top MTBE layer enriched with lipids was taken and dried under nitrogen gas before mixing with 3 mL Isopropanol and 3 mL methanol to make Resuspension Solvent. The extract was resuspended in 300 µl Resuspension Solvent, followed by centrifugation at 13,000 × g for 10 min. 200 µl on the top was collected and then transferred into the glass autosampler tubes. Phospholipid classes were solved on the state-of-the-art Orbitrap Q Exactive Mass Spectrometer (Thermo Scientific) coupled with HILIC and reversed-phase liquid chromatography. Quantification of individual molecular phospholipid species of PE or PS was calculated by determining the ratio of the peak area of each molecular phospholipid species to the corresponding internal SPLASH standard. The fold change of total mitochondrial PE or PS species and the relative abundance (mol%) of individual PE or PS species in the PE or PS class of mitochondria were analyzed, respectively. The species of each phospholipid head group were summed together to obtain the total abundance of each head group which was used to calculate the percentage of relative abundance of PE or PS. As for the heat map construction, the values of individual molecular phospholipid species of PE or PS were normalized by the internal SPLASH standard. Z-score was calculated for each molecular phospholipid species in non-Tg or PS19 groups using the formula z = (x-μ)/σ, where x was the raw score, μ was the population mean, and σ was the population standard error of the mean. Double gradient heat map was then generated for these significant differential phospholipid species using the Z-Score with GraphPad Prism 10. Comparisons of lipidomics studies were performed by one-way ANOVA with Bonferroni’s correction (compared to the control group), as previously described [117-119].

### Tissue preparation and immunohistochemistry

Animals were anesthetized with 2.5% avertin (0.35ml per mouse) and transcardially perfused with fixation buffer (4% paraformaldehyde in PBS, pH 7.4). Brains were dissected out and postfixed in fixation buffer overnight and then placed in 30% sucrose at 4°C. 10-µm-thick coronal sections were collected consecutively to the level of the hippocampus and used to study the co-localization of various markers. After incubation with blocking buffer (5% goat serum, 0.3% Triton X-100, 3% BSA, 1% glycine in PBS) at RT for 1 hr, the sections were incubated with primary antibodies at 4°C overnight, followed by incubation with secondary fluorescence antibodies at 1:600 dilution at RT for 1 hr. After fluorescence immunolabeling, the sections were stained with DAPI and washed three times in PBS. The sections were then mounted with an anti-fading medium (vector laboratories, Cat# H-5000) for imaging. Confocal images were obtained using an Olympus FV3000 oil immersion 40× objective with a sequential-acquisition setting. Eight to ten sections were taken from top-to-bottom of the specimen and brightest point projections were made. Fluorescence intensities of LC3, AT8, PHF1, synaptophysin (SYP), NeuN, LAMP1, or p62/SQSTM1 were expressed in arbitrary units of fluorescence per square area. The mean intensities of LC3, AT8, PHF1, SYP, NeuN, or LAMP1 in the hippocampi PS19 mouse brains and/or rTg4510 were normalized to those in the brains of non-Tg littermate, PS19, or rTg4510 mice without glutamine supplementation. As for quantification of co-localization to define the densities of autolysosomes co-labeled by LC3 and LAMP1, a threshold intensity was preset for both fluorescent signals, which was determined with the thresholding function of NIH ImageJ described [20,113,120]. The co-localized pixels above the threshold intensity were automatically quantified and scored by ImageJ based on the fluorescence intensity profile, which was expressed as co-localized mean intensity positive for both channels. Co-localization was presented as co-localized mean intensities of LC3 with LAMP1, which were normalized to those from control non-Tg littermate, PS19, or rTg4510 mice fed with regular water. Data were obtained from two to three pairs of PS19 mice or rTg4510 mice and the number of imaging brain slices used for quantification was indicated in the figures.

### Transmission electron microscopy

Hippocampi from 6-month-old PS19 mouse brains with and without 3-month glutamine administration were cut into small specimens (one dimension < 1 mm) and fixed in Trump’s fixative (Electron Microscopy Sciences) for 2 hr at RT. The sections were then washed with 0.1 M Cacodylate buffer and postfixed in 1% osmium tetroxide, followed by dehydrating in ethanol and embedding using the EM bed 812 kit (Electron Microscopy Sciences) according to a standard procedure. Images were acquired on an electron microscope (1200EX; JEOL) (Electron Imaging Facility in the Department of Pathology and Laboratory Medicine, Robert Wood Johnson Medical School). For quantitative studies, the number of autophagic vacuole (AV)-like organelles and the number of mitochondria at presynaptic terminals were counted from electron micrographs. AVs were characterized by double-membrane structures along with other organelles or vesicles, or containing partially degraded cytoplasmic material with higher electron density [109,121,122]. Quantification analysis was performed blindly to condition.

### Behavior assays

Male mice only were assessed for cognitive and emotional behavior. Groups contained 7-12 male mice/group for each behavioral task. Tests were carried out in mice at 8 months of age with and without 5-month glutamine supplementation. All tests used the ANY-maze Video Tracking System (Stoelting Co.) with additional video review by experimenters who were blind to treatment group conditions.

Open field test: This test involved the measurement of spontaneous locomotor activity and within-session habituation to a novel environment as previously described [81]. The apparatus was made of black Plexiglas sheets (56 × 62 × 28 cm) to form an open box with high walls. The inner bottom and inner walls of the apparatus were painted in blue, with the floor evenly divided by 5 × 6 red lines into thirty identical cells. Mice were allowed to explore the open field for 5 min. Total movements (ambulations) were tracked using ANY-maze, which was set to track movements in an outer periphery area and center (inner) region of the open field, away from the periphery. The apparatus was thoroughly cleaned with 70% ethanol between each animal.

Sociability and preference for social novelty test: The test for sociability and preference for social novelty was conducted as previously described [72,77-79]. The apparatus consisted of a rectangular, three-chambered box. Each chamber was measured 26.5 × 46 × 30 cm with the dividing walls made of white Plexiglas and with small square openings (4 × 4 cm) allowing access into each chamber. The stranger mouse (stranger 1 (S1), a 7-month-old male C57BL/6 mouse that was not a littermate nor previously used in any other experiments) was enclosed in a small, circular metal wire cage (10 × 10 cm) that allowed nose contact between the bars but prevented fighting. For the first 10-min session, the placement of S1 in the left or right-side chamber was systematically alternated between trials, with the opposite side chamber left empty (E). The test mouse was first placed in the middle chamber and allowed to explore the entire social test box. Using the ANY-maze Video Tracking System (Stoelting Co.), the movement of the test mouse and the amount of time that the test mouse spent in each chamber were tracked for 10 min for subsequent data analysis to quantify social preference for S1. After the sociability test session was over, there was a 10-min inter-trial interval before the test mouse was guided to the center chamber for a social recognition test in a second 10-min session. Also, prior to this session, a second, unfamiliar male mouse (stranger 2 (S2), 7 months old) was introduced into the empty side chamber. This S2 mouse was enclosed in an identical small wire cage to the S1 mouse. The test mouse had a choice between the first, already-investigated familiar mouse (S1) and the novel unfamiliar mouse (S2). As described above, the amount of time spent in each chamber with S1 or S2 during this second 10-min session was recorded for the measurement of social recognition memory. Thus, sociability and recognition memory were assessed in trials by analyzing the amount of time spent exploring the empty chamber E or the chamber with S1 in the first session and the chambers containing S1 or S2 in the second session, respectively. The stranger mice used in this experiment were 7-month-old C57BL/6J male mice, not littermates.

Morris water maze: The water maze consisted of a pool (110 cm in diameter) containing tap water (24°C ± 1°C) made opaque with non-toxic powdered paint. A platform (9.4 cm in diameter) was submerged 1.5 cm beneath the water surface. The pool was surrounded by a curtain and a facing wall. Distinct cues were placed on the curtain and wall and remained the same for all animals. Mice were trained to locate the hidden platform in a single day [81], with a probe memory trial given after the final acquisition trial. Overall training was conducted in four blocks. The first block (Block 1) of trials consisted of three trials in which the target platform was made visible by an affixed flag. These three flagged trials ensured that animals were primed to seek an escape platform, as well as tested for any visual deficits. Blocks 2-4 consisted of hidden-platform training, with each block consisting of four trials, with the platform hidden below the surface, off-center and randomly located (across animals) in one of the four quadrants of the circular pool. For each training block, the inter-trial interval was 5 min, such that each mouse completed a block in approximately 20 minutes. For each trial in a block, mice were placed into the water with the head facing the tank wall and allowed a maximum of 60 sec to find the hidden platform (mice were returned to their holding cage until the next trial if they had not found the platform by 60 seconds). Each succeeding trial within a block involved a different starting location (north, south, west, or east relative to the platform). Once a block was completed, it was repeated for each mouse, such that all mice were run through three hidden-platform training blocks (Blocks 2-4), which consisted of a total of 12 acquisition trials. 1 hour or 24 hours after the last trial of Block 4, a memory 60-second probe trial was conducted by removing the platform and allowing mice to search for the platform. Data consisted of the time to locate the hidden platform during learning, and the time spent in the target quadrant, as well as path length, during the probe test. These data were calculated by the ANY-maze Video Tracking System (Stoelting Co.).

Contextual fear conditioning: Conditioning was conducted in Coulbourn operant learning chambers that measured 19 × 20.3 × 30.5 cm and were located inside sound-attenuating cabinets (Coulbourn Instruments, Whitehall, PA). The chamber contained a steel grid floor connected to a programmable shocker, while the walls were made of clear Plexiglas. For contextual fear conditioning, the mice were placed within the conditioning chamber for 3 min to develop a representation of the context prior to the onset of a single unconditioned stimulus (US; 1.0 mA footshock; 1 second duration). Following the shock, they were allowed to remain in the chamber for 2 min, during which immediate freezing was measured continuously. Mice were then returned to the home cages. Memory was tested 24 hours after training for 4 min in the same conditioning chamber. Animal movements were tracked with the ANY-maze Video Tracking System (Stoelting Co.).

### Statistical analysis

Statistical parameters including the definitions and exact value of n (e.g., number of neurons, number of neuronal soma, number of axons, number of synapses, number of mitochondria, number of mouse brain slice sections, number of animals, etc), deviations, p values, and the types of the statistical tests were reported in the Figures and in the corresponding Figure Legends. p values from one-way analysis of variance (ANOVA) or Kruskal-Wallis test for the analysis of the data from three or more groups were indicated in the corresponding text. Statistical analysis was carried out using GraphPad Prism (version 10.1.1). All statistical comparisons were conducted on data originating from three or more biologically independent experimental replicates. Statistical comparisons between the two groups were performed by an unpaired student’s *t*-test (sample size ≥ 15) or a Mann-Whitney U test (sample size < 15). Comparisons between three or more groups were performed by one-way analysis of variance (ANOVA) with post hoc testing by Bonferroni’s correction (sample size ≥ 15) or by Kruskal-Wallis test with Dun’s multiple comparison post hoc test or Bonferroni’s correction (sample size < 15) where otherwise indicated. Data are expressed as mean ± SEM. Differences were considered significant with p < 0.05.

## Acknowledgments

We are grateful to E. Gavin, G. Rajapaksha, K. Vandersleen, S. Agarwal, and other members of the Cai Lab for their research assistance and constructive discussion; Y. Zuo and J. Cheung for critical reading; X. Su at the Metabolomics Core Facility of Rutgers Cancer Institute of New Jersey; R. Patel at the EM facility in the Department of Pathology and Laboratory Medicine, Robert Wood Johnson Medical School for technical help; H. Imamura for AT1.03 and mitAT1.03 plasmids. P. Davies for PHF1 antibody. This research was supported by National Institutes of Health grants R01NS089737 and R01GM135326 (Q.C.).

