## Supplemental information for "Mitochondrial bioenergetics stimulates autophagy for pathological tau clearance in tauopathy neurons"

### Figure S1 (Cai)

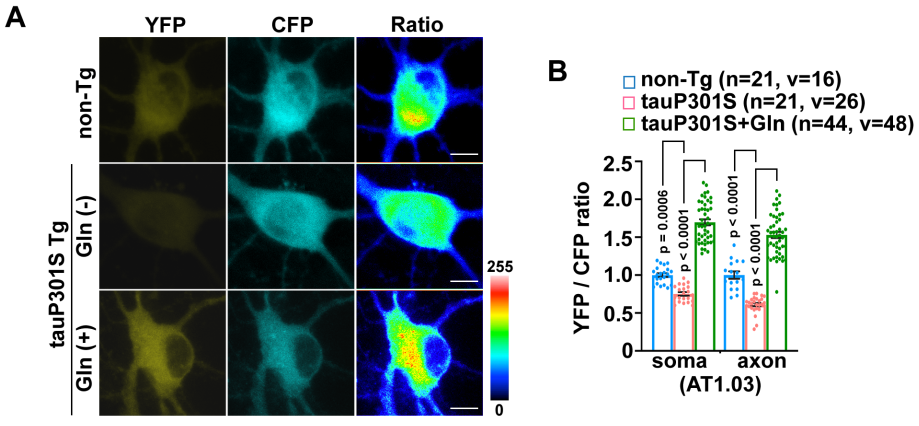


**Figure S1. Anaplerotic stimulation corrects energy dysfunction and synaptic mitochondrial bioenergetic deficits in tauopathy neurons.** (A-B) Representative images (A) and quantitative analysis (B) of AT1.03-indicated cellular ATP levels in DIV10-12 primary cortical neurons derived from non-transgenic (Tg) and tauP301S Tg (PS19) mouse brains with and without 10 mM glutamine treatment for 24 hours. The YFP/CFP ratios in the soma and the axons of PS19 neurons in the presence and absence of glutamine were normalized to those in control non-Tg neurons. Gln: glutamine. Data were collected from the total number of neuronal soma (n) and axons (v) as indicated in parentheses (B) from three independent experiments. Data were expressed as the mean ± SEM with dots as individual values and analyzed by one-way ANOVA with Bonferroni's correction. Scale bars: 10 μm.

### Figure S2 (Cai)

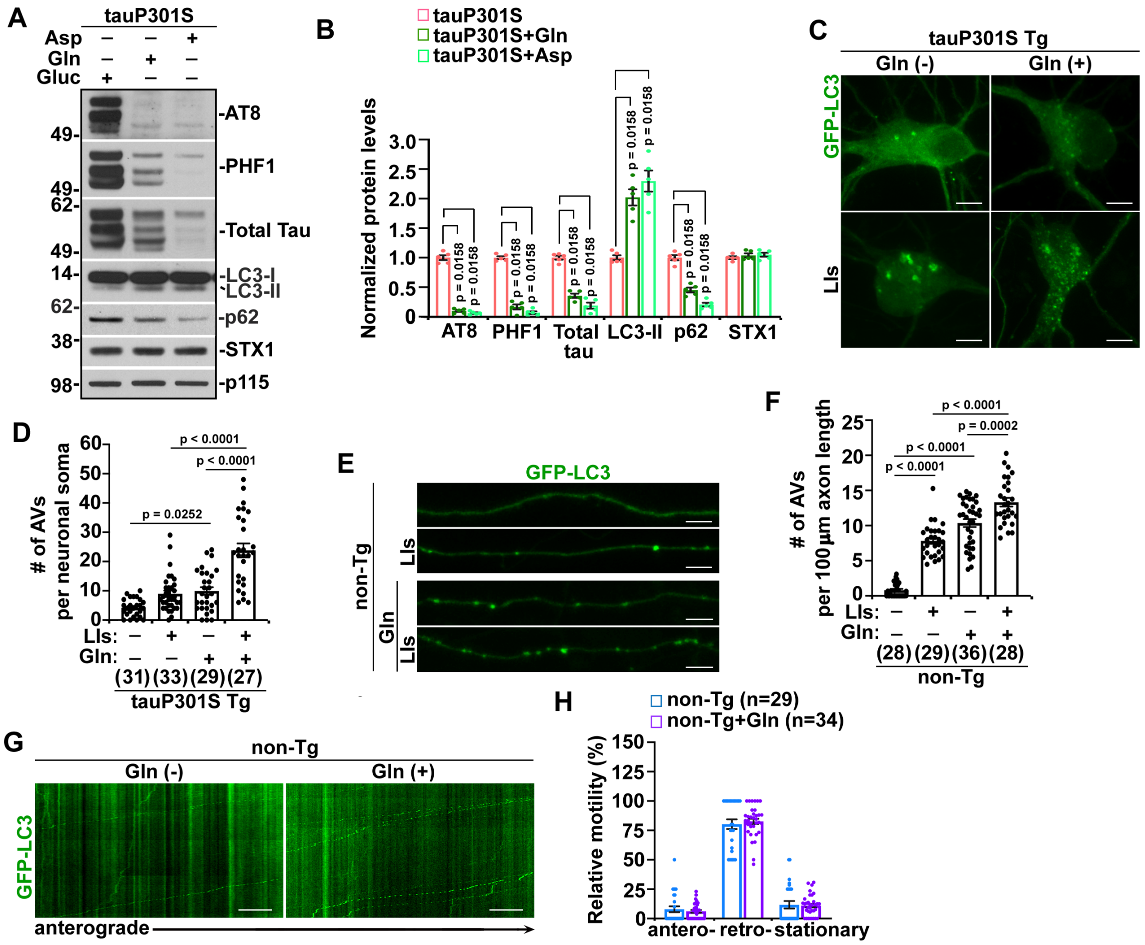


**Figure S2. Anaplerosis-boosted OXPHOS leads to increased AV biogenesis and autophagic clearance in non-Tg and PS19 neurons.** (A-B) Representative blots (A) and quantitative analysis (B) of autophagy and tau protein levels in DIV12-14 PS19 neurons incubated for 24 hours with glucose (25 mM), glutamine (10 mM), or aspartic acid (5 mM). The protein levels were normalized to the loading control p115 and to those of PS19 neurons with glucose, respectively. Gluc: glucose; Asp: aspartic acid; STX1: Syntaxin 1. p115, a Golgi protein. (C-D) Representative images (C) and quantitative analysis (D) of GFP-LC3-labeled AV density in the soma of DIV10-12 PS19 neurons in the presence and absence of LIs (20 µM E64D and 20 µM Pepstatin A), glutamine, or glutamine and LIs. The data were quantified and expressed as the number of GFP-LC3-marked AVs per neuronal soma. AV: autophagosome/autophagic vacuole; LIs: lysosomal inhibitors. (E-F) Representative images (E) and quantitative analysis (F) of autophagic flux in DIV10-12 non-Tg axons in the presence and absence of LIs, glutamine, or glutamine and LIs. The data were quantified and expressed as the number of GFP-LC3-marked AVs per 100 µm axonal length in non-Tg neurons. (G-H)

Representative kymographs (I) and quantitative analysis (H) of relative motility of AVs in the axons of non-Tg neurons with and without 24-hour glutamine incubation. Data were quantified from five independent repeats (B) or were collected from the total number of neurons (n) as indicated in parentheses (D, F, and H) from at least three independent experiments. Data were expressed as the mean ± SEM and analyzed by Kruskal-Wallis test with Bonferroni's correction (B), one-way ANOVA with Bonferroni's correction (D and F), or two-sided unpaired Student's *t*-test (H). Scale bars: 10 μm.

**Figure S3 (Cai)**

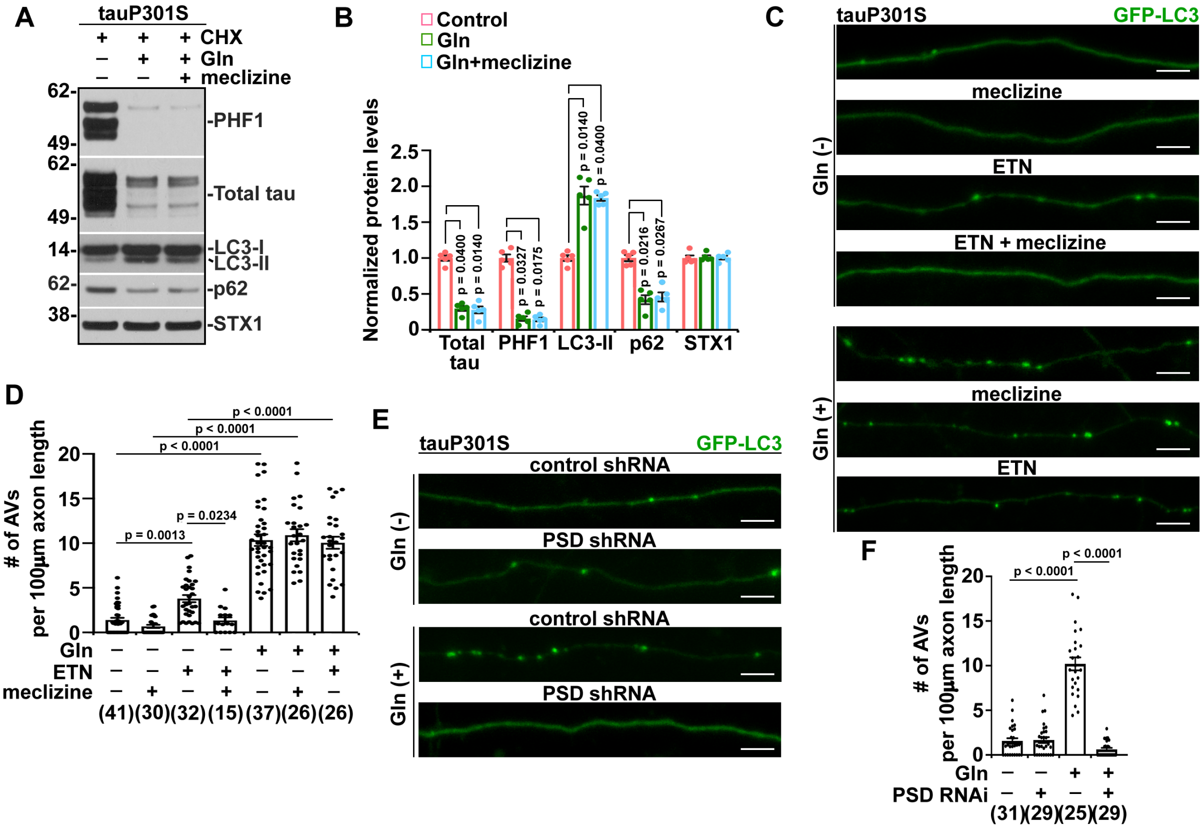

**Figure S3. Anaplerotic metabolism-stimulated autophagy in tauopathy neurons is independent of phosphatidylethanolamine (PE) biosynthesis in the endoplasmic reticulum (ER).** (A-B) Representative blots (A) and quantitative analysis (B) of autophagy and tau protein levels in DIV12-14 PS19 neurons treated for 24 hours with and without 10 mM glutamine or 10 mM glutamine and 5 μM meclizine in the presence of 10 μg/ml CHX. The protein levels were normalized to STX1 and to those of control PS19 neurons with CHX alone, respectively. (C-D) Representative images (C) and quantitative analysis (D) of AV density in DIV10-12 PS19 axons treated for 24 hours with vehicle, 5 μM meclizine, 5 mM ETN, both, or glutamine with and without 5 μM meclizine or 5 mM ETN. The data were quantified and expressed as the number of AVs per 100 μm axonal length in PS19 neurons. ETN: ethanolamine. (E-F) Representative images (E) and quantitative analysis (F) of autophagic flux in DIV10-12 PS19 axons expressing control or PSD shRNA in the presence and absence of 10 mM glutamine. The data were quantified and expressed as the number of GFP-LC3-labeled AVs per 100 μm axonal length in PS19 neurons. PSD: phosphatidylserine decarboxylase. Data were quantified from five independent repeats (B) or were collected from the total number of neurons (n) as indicated in parentheses (D and F) in three independent experiments. Data were expressed as the mean ± SEM and analyzed by Kruskal-Wallis test with Dun’s multiple comparison post hoc test (B), one-way ANOVA with Bonferroni’s correction (D and F). Scale bars: 10 μm.

### Figure S4 (Cai)

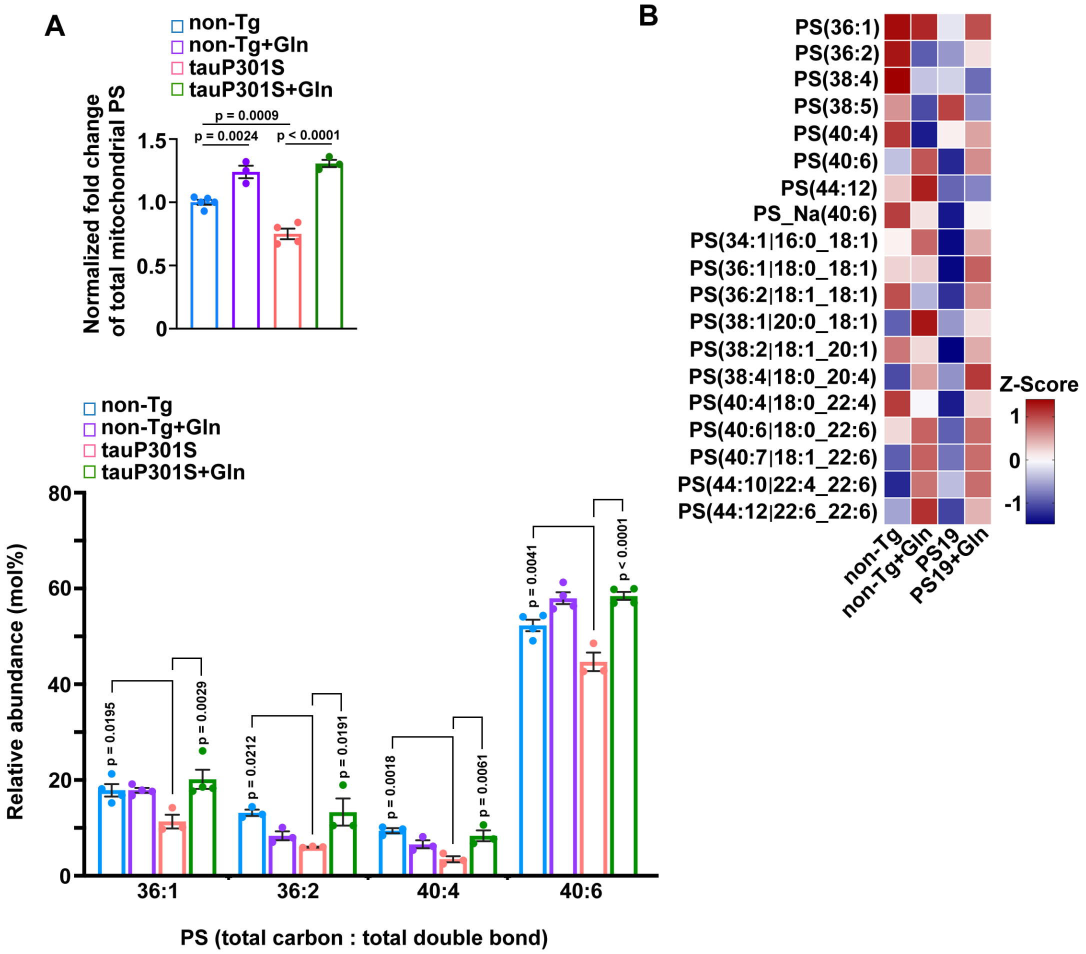


**Figure S4. Increased PS levels in the brain mitochondria of tauopathy mice with glutamine supplementation.** (A) Lipidomics analysis of the fold change of total mitochondrial PS species in the brains of 6-month-old non-Tg and PS19 mice with and without glutamine supplementation. The changes of mitochondrial PS in the brains of PS19 control mice or non-Tg and PS19 mice treated with glutamine were normalized to that of control non-Tg mice fed with water. (B) Heat map analysis for molecular phospholipid species of PS extracted from 6-month-old non-Tg and PS19 mouse brain mitochondria. Rows represent the mean values for PSs and columns indicate those of non-Tg or PS19 mice administered with water or glutamine. (C) Relative abundance (mol%) of individual PS species in the total PS class of mitochondria in non-Tg and PS19 mouse brains administered with water or glutamine. Data were quantified from three to four animals (A-C) in each group. Data were expressed as the mean ± SEM and analyzed by one-way ANOVA with Bonferroni's correction (A and C).

**Figure S5 (Cai)**

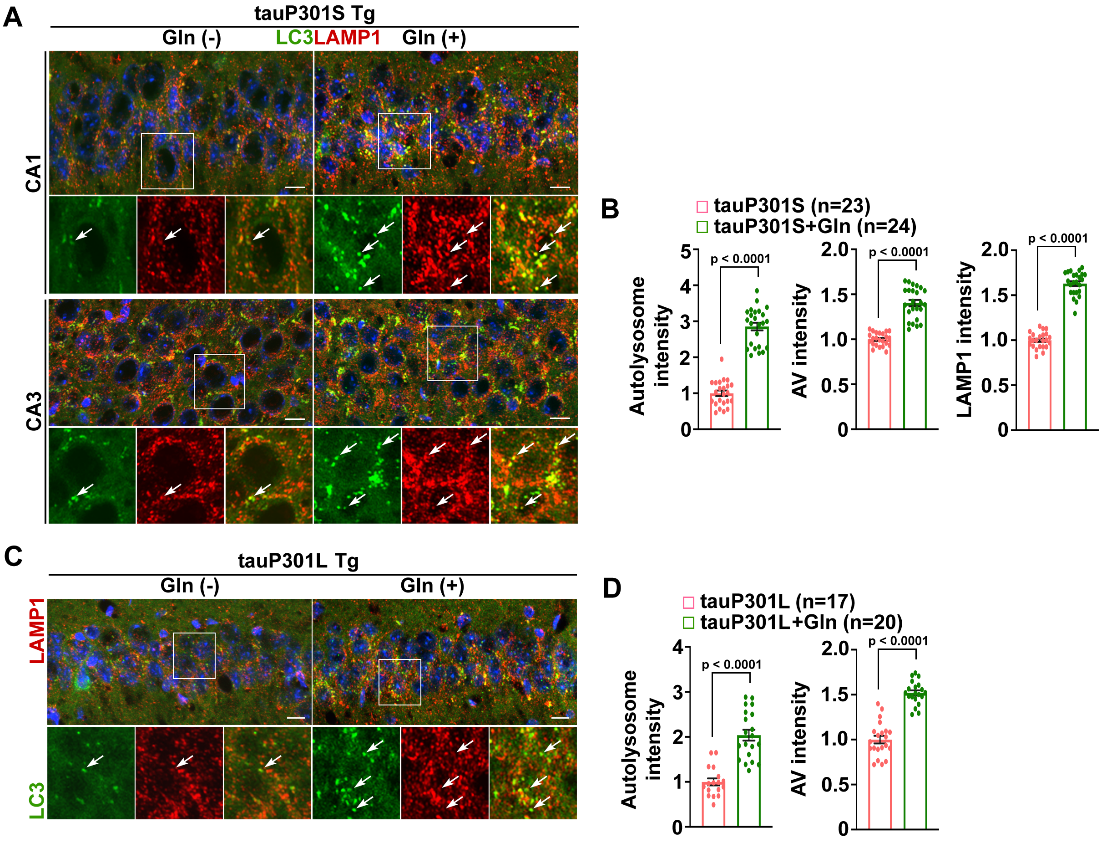


**Figure S5. Elevated autophagy activity in glutamine-supplemented tauopathy mouse brains.** (A-B) Representative images (A) and quantitative analysis (B) of autophagy activity in the soma of hippocampal neurons in 9-month-old PS19 mouse brains with and without glutamine supplementation. Data were expressed as the densities of autolysosomes (arrows), AVs, or LAMP1-marked autolysosomes/lysosomes in the neuronal soma of hippocampal CA3 regions in PS19 mouse brains and quantified from a total number of imaging slice sections as indicated in parentheses (B) from three pairs of PS19 mice. (C-D) Representative images (C) and quantitative analysis (D) of AVs and autolysosomes (arrows) in the soma of hippocampal neurons in 6-month-old rTg4510 mouse brains with and without glutamine supplementation. Data were expressed as the densities of autolysosomes or AVs in the neuronal soma of hippocampal CA1 regions in rTg4510 mouse brains and quantified from a total number of imaging slice sections as indicated in parentheses (D) from two rTg4510 mice in each group. Data were expressed as the mean ± SEM and analyzed by two-sided unpaired Student's *t*-test (B and D). Scale bars: 10 μm.

**Figure S6 (Cai)**

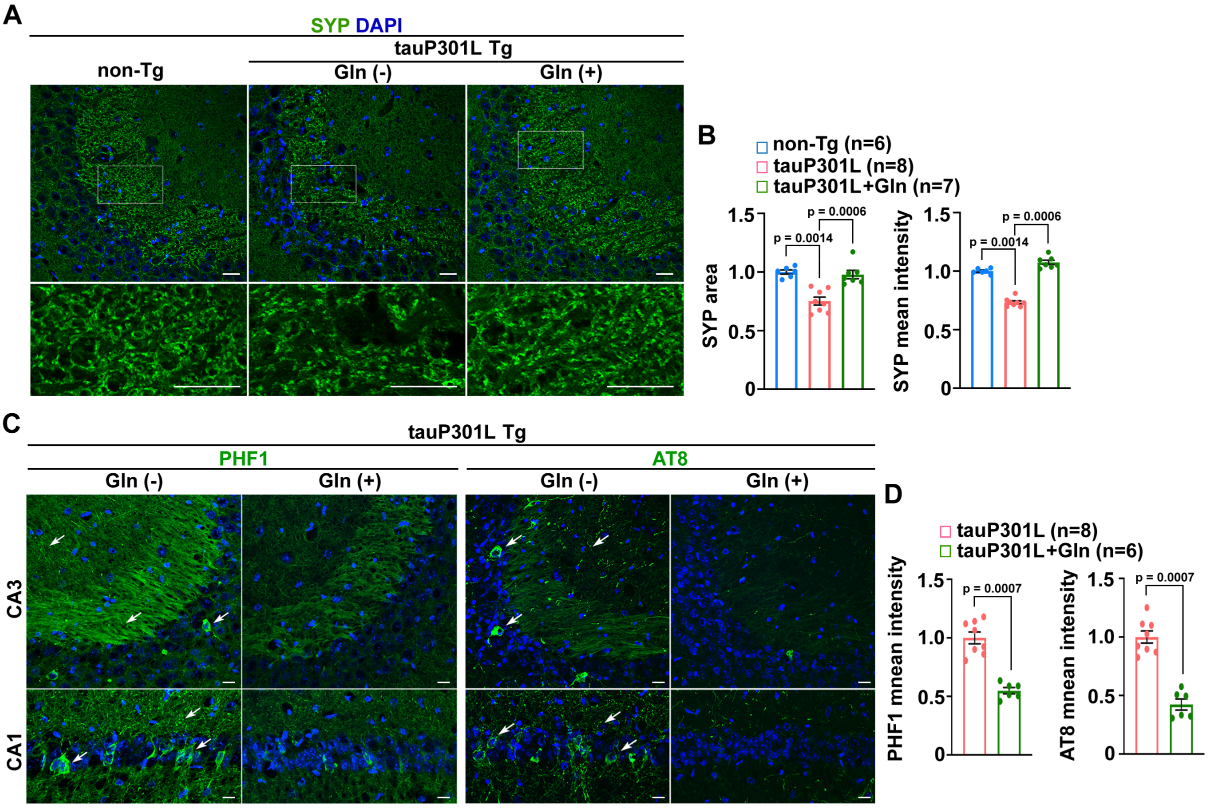


**Figure S6. Attenuations of synaptic defects and tau pathology in the brains of tauopathy mice under glutamine metabolism.** (A-B) Representative images (A) and quantitative analysis (A) of presynaptic density in the hippocampal mossy fibers of 6-month-old non-Tg mice and rTg4510 mice administrated with water or glutamine. The area and the mean intensity of SYP-marked presynaptic terminals were quantified and normalized to those of control non-Tg littermates, respectively (F). Data were quantified from a total number of imaging slice sections indicated in parentheses (F) from two mice in each group. (E-F) Representative images (E) and quantitative analysis (F) of phospho-tau levels in the hippocampal regions of 6-month-old rTg4510 mouse brains with and without 16-week glutamine treatment. The mean intensities of PHF1 and AT8 antibody-labeled phospho-tau structures (arrows) were quantified and normalized to control rTg4510 mice fed with water from a total number of imaging slice sections indicated in parentheses (D) from two pairs of mice. Data were expressed as the mean ± SEM and analyzed by Mann-Whitney U test (B) or Kruskal-Wallis test with Bonferroni's correction (D). Scale bars: 10 μm (A and C), and 25 μm (E).

### Figure S7 (Cai)

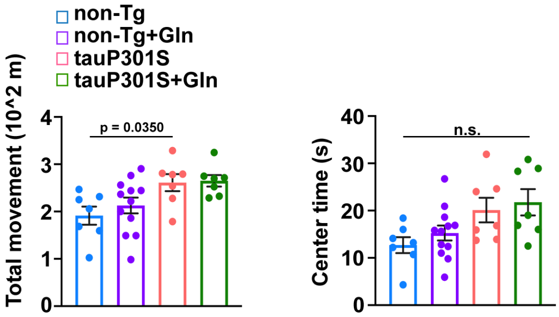


**Figure S7. Open field test in bioenergetically enhanced PS19 mice.** Open field test performed in non-Tg and PS19 male mice with and without glutamine supplementation (n = 7-12 mice per group; 8 months of age). Data were shown as the mean ± SEM and analyzed by Kruskal-Wallis one-way analysis of variance test with Bonferroni's correction. n.s.: non-significant.
